# Mitochondrial cytochrome *c* liberates the nucleophosmin-sequestered ARF tumor suppressor in the nucleolus

**DOI:** 10.1101/2020.05.07.057075

**Authors:** Katiuska González-Arzola, Antonio Díaz-Quintana, Noelia Bernardo-García, Miguel Á. Casado-Combreras, Carlos A. Elena-Real, Alejandro Velázquez-Cruz, Sergio Gil-Caballero, Adrián Velázquez-Campoy, Elzbieta Szulc, Isabel Ayala, Rocío Arranz, Xavier Salvatella, José M. Valpuesta, Juan A. Hermoso, Miguel A. De la Rosa, Irene Díaz-Moreno

## Abstract

The alternative reading frame (ARF) protein is crucial in the cellular response to oncogenic stress, being likewise the second most frequently inactivated gene in a wide spectrum of human cancers. ARF is usually sequestered in the nucleolus by the well-known oncogenic nucleophosmin (NPM) protein and is liberated in response to cell damage to exhibit its tumor-suppressor ability. However, the mechanism underlying ARF activation is unknown. Here we show that mitochondria-to-nucleus translocation of cytochrome *c* upon DNA damage leads to the break-off of the NPM-ARF ensemble and subsequent release of ARF from the nucleoli. Our structural and subcellular data support a molecular model in which the hemeprotein triggers the extended-to-compact conformation of NPM and competes with ARF for binding to NPM.

## Main Text

The alternative reading frame (ARF) protein of the INK4a-ARF gene constitutes a pivotal stress signaling and response pathway (*1*). Stress raises DNA damage levels in cells, in turn activating the ARF/p53 route, which initiates adaptive responses (*2*). INK4a-ARF is the second most frequently inactivated gene in human cancers (*3*). In mammalians, ARF (p19ARF in mice, p14ARF in humans) regulates p53 levels by inactivating the p53 ubiquitin ligase HDM2 (human double minute clone 2, MDM2 in mice) (*4*).

ARF is however usually sequestered in the nucleolus by nucleophosmin (NPM) – a.k.a. B23, NO38 or numatrin. NPM (*5*) is a histone chaperone that participates in ribosome synthesis (*6*), cytoplasmic-nuclear shuttling (*7*), chromatin remodeling (*8*) and mitotic spindle assembly (*9*). NPM is also a key factor favoring DNA repair and promoting cell survival (*10*). Expectedly, NPM mutations or dysregulation often correlate with oncogenesis (*5,10*). Under homeostasis, NPM sequesters ARF in the nucleolus, thus enabling p53 ubiquitination and subsequent proteasomal degradation (*3,11,12*). Upon stress and/or DNA damage, NPM releases ARF, which then binds to HDM2 to diminish p53 degradation (*10*). In fact, cells enter apoptosis after knocking down NPM (*13*). However, the molecular mechanism triggering NPM dissociation from ARF remains unknown.

In this work, NPM emerges as a novel target of mitochondrial cytochrome *c* (C*c*) in human cells undergoing DNA damage. A recent proteomic analysis showed that C*c* targets several histone chaperones upon DNA damage (*14,15*). Indeed, C*c* moves into the cell nucleus and binds SET/TAF-Iβ once DNA lesions are detectable without stimulation of death receptors or stress-induced pathways (*16*). In plants, C*c* likewise targets histone chaperone NRP1, analogue to human SET/TAF-Iβ, under similar conditions (*17,18*).

Here, we found that mitochondrial endogenous C*c* is translocated into the nucleus of tumor and non-tumor cells following DNA breaks. Further, C*c* competes with ARF for binding to the N-terminal oligomerization domain of NPM. To understand the molecular basis of such competition, we determined the structure of the complex between NPM and C*c*. Notably, C*c* lodges in the cavity formed by the NPM arms, the acidic disordered middle region of NPM, stabilizing the NPM-C*c* complex. Our biophysical and structural analyses reveal that a single molecule of C*c* can release up to five molecules of ARF out of NPM. Further liquid-liquid phase separation assays demonstrate that NPM undergoes phase transitions via heterotypic interactions with C*c*, as does NPM with ARF. Noteworthy, C*c* contains lysine-rich motifs rather than the canonical, linear arginine-rich motifs of well-known membrane-less organelle components, such as ARF, and forms nucleolar-like droplets with NPM. Even more interesting is the fact that C*c* can enter the nucleoli to break off the preformed NPM-ARF ensembles and induce ARF release, thereby explaining the molecular mechanism underlying ARF activation upon DNA damage.

### Nucleophosmin as a nuclear target of cytochrome *c* upon DNA breaks

First, to ascertain whether mitochondrial C*c* can reach nuclei upon DNA damage, we immunodetected endogenous C*c* in cells treated with either camptothecin (CPT) or doxorubicin. Immunostaining of HeLa cells treated with these drugs revealed the release of C*c* from mitochondria to the cytoplasm and the nucleus (Fig. 1a, *upper*). Mitochondria were labeled with mitotracker—whose accumulation depends on mitochondrial transmembrane potential—because C*c* release from mitochondria precedes the loss of such potential, as previously described (*19*) (Fig. 1a and Fig. S1a). We further explored the C*c* subcellular location in non-cancer cells treated similarly. We observed a parallel behavior in CV-1 in origin, carrying SV40 cells (Cos-7, Fig. 1a, *lower*), as well as in mouse embryonic fibroblasts (MEF) (Fig. S1a, *left*). These results were also evident when applying an alternative immunofluorescence method in HeLa cells (Fig. S1a, *right*). In a previous report (*16*), we performed subcellular fractioning assays to demonstrate that endogenous C*c* translocates into the nuclei upon DNA breaks induced by CPT, doxorubicin or indotecan. On the contrary, C*c* was unable to be translocated into cell nuclei upon treatment with other apoptosis-inducing agents which do not induce DNA damage, namely tumor necrosis factor (TNF)-related apoptosis-inducing ligand (TRAIL) and staurosporine (STP) (*16*). More interestingly, we showed that C*c* translocation into nucleus in response to DNA breaks occurs prior to caspase-3 activation (*16*).

**Fig. 1.**
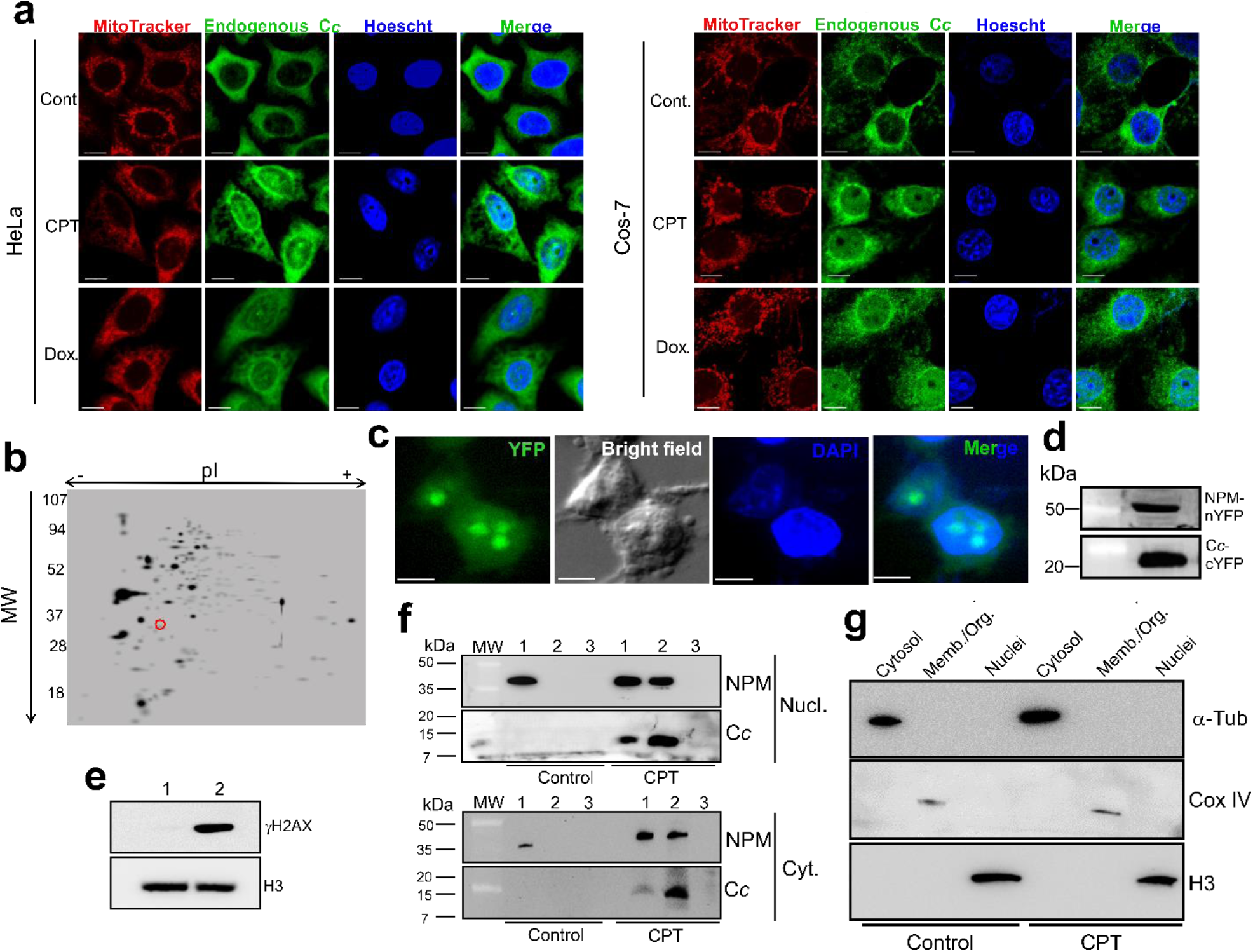
DNA damage-induced nuclear translocation of C*c* and identification of NPM as a C*c* nucleo-cytoplasmic target. **a**, Immunofluorescence analysis of endogenous C*c* in HeLa (*upper*) and Cos-7 (*lower*) cells, upon treatment with 20 μM CPT or 4 μM doxorubicin for 6 h. Subcellular distribution of C*c* was visualized with an anti-C*c* antibody (green fluorescence) using a confocal microscope (x63 oil objective). Mitochondria were stained with MitoTracker Red CMXRos (red fluorescence) and nuclei with Hoechst (blue). Non-treated cells were used as control. Co-localization of green C*c* fluorescence and blue nuclear staining is shown in the merged images. Scale bars are 10 μm. HeLa cells were fixed in a 95% ethanol and 5% acetic acid solution, while Cos-7 cells were fixed in 2% PAF. **b**, 2D SDS-PAGE gel of Jurkat T cell extracts sieved by affinity towards E104C C*c*. Red circle highlights NPM. **c-d**, BiFC and western blot analyses showing the *in-cell* interaction. HEK293T cells were transfected with the N-terminal YFP fragment attached to NPM (NPM-nYFP), along with a vector containing the C-end domain of YFP bound to C*c* (C*c*-cYFP). Images were taken 24 h after transfection and 6 h of 10 μM CPT treatment (**c**). Scale bars are 5 μm. Protein expression was checked by immunoblotting *vs*. anti-EGFP antibody (**d**). **e**, γ-H2AX detection in the nuclear lysates of HeLa cells upon treatment with 20 μM CPT for 4 h. Lanes 1 and 2 are control and CPT-treated samples, respectively. **f**, IP of endogenous NPM with C*c* in the nucleus (*upper*) and cytosol (*lower*) of HeLa cells treated with 20 μM CPT for 4 h. Western blot detection of NPM (∼35 kDa) in non-treated and treated cells, along with C*c*-IP of lysates from non-treated and CPT-treated cells probed with the NPM antibody. Lanes 1 and 2 are lysate and C*c* IP, respectively. Mouse immunoglobulin G (IgG) was used as control (lane 3). Detection of C*c* as a ∼15 kDa band in raw and C*c*-immunoprecipitated nuclear lysates are also shown following CPT treatment. **g**, Purity of subcellular fractions of non-treated and CPT-treated HeLa cells was verified by immunoblotting against α-Tub (55 kDa), COX-IV (14 kDa) and histone H3 (14 kDa) antibodies for detecting cytosolic, membrane and nuclear proteins, respectively. Cells were treated with 20 μM CPT for 4 h.

We extended our previous proteomic analysis of the C*c*-centered interactome—based on tandem affinity chromatography, 2D-PAGE and mass spectrometry (*14*). A C-end cysteine-substituted C*c* (E104C) tethered to a thiol-Sepharose 4B matrix allowed sifting extracts from Jurkat cells either untreated or treated for 6 h with 10 μM camptothecin (CPT), which primarily induces single strand breaks that can further be converted to DNA double strand breaks (DSBs) (*20*). 2D-PAGE experiments enabled us to detect up to 21 putative targets, accessible to C*c* upon DNA damage but not under homeostasis. Mass spectrometry analysis identified the spot marked in Fig. 1b in red as NPM.

Bimolecular fluorescence complementation (BiFC) experiments (*21*) confirmed the interaction in cells. A construct of C*c* fused to the C-end domain of the yellow fluorescent protein (YFP) was available at our lab (*14*), and we built another one encoding full length NPM (hereafter NPM_(1-294)_) ahead of the N-end fragment of the YFP (nYFP). After co-transfection, 10 μM CPT was added to the culture. The fluorescence pattern and DAPI overlay (Fig. 1c) clearly show that NPM interacts with C*c* in both the nucleus and the cytoplasm. Highly fluorescent spots within nuclei pinpoint condensates of NPM molecules. Protein expression was confirmed by Western blot analysis (Fig. 1d). Control cells co-transfected with bFos and bJun chimeras displayed YFP reconstitution. They did not when using bFosΔZip construct lacking the Jun-binding domain (Fig. S1b).

C*c* migrates into the cell nucleus upon DSBs accumulation solely (*16*). We confirmed the DNA damage after 4 h treatment by probing the accumulation of H2AX (γH2AX) (Fig 1e). Accordingly, phosphorylated histone H2AX (γH2AX), which quickly accrues after DNA damage, is detected in cell lysates 4 h after treatment (Fig 1e). Moreover, we detected endogenous NPM in Western blots of C*c* immunoprecipitated cell extracts from HeLa cell line. Co-immunoprecipitation of proteins was observed from both cytoplasmic and nuclear fractions only upon DNA damage, 4 h after adding 20 μM CPT (Fig. 1f), but not from the lysates of untreated cells. When using IgG for IP, none of the NPM-featuring band was detected (Fig. 1f). Immunoblotting against α-tubulin, COX-IV (cytochrome *c* oxidase subunit IV) and histone H3 antibodies confirmed the purity of subcellular fractions (Fig. 1g).

### Cytochrome *c* competes with p19ARF for NPM binding

NPM participates in DNA damage response triggered by radiation (*10*). Indeed, NPM modulates the interaction between two p53 regulators –HDM2 and ARF– (*12, 13*) (Fig. 2a). We hypothesize that C*c* plays a role in ARF release form the NPM complex upon DNA damage by competition for the binding site in NPM. To understand the molecular details of both interactions and test the hypothesis we designed several NPM constructs and assessed their ability to bind p19ARF and C*c* (Fig. 2b) *in vitro*. All NPM constructs folded correctly (Fig. S2a-d).

**Fig. 2.**
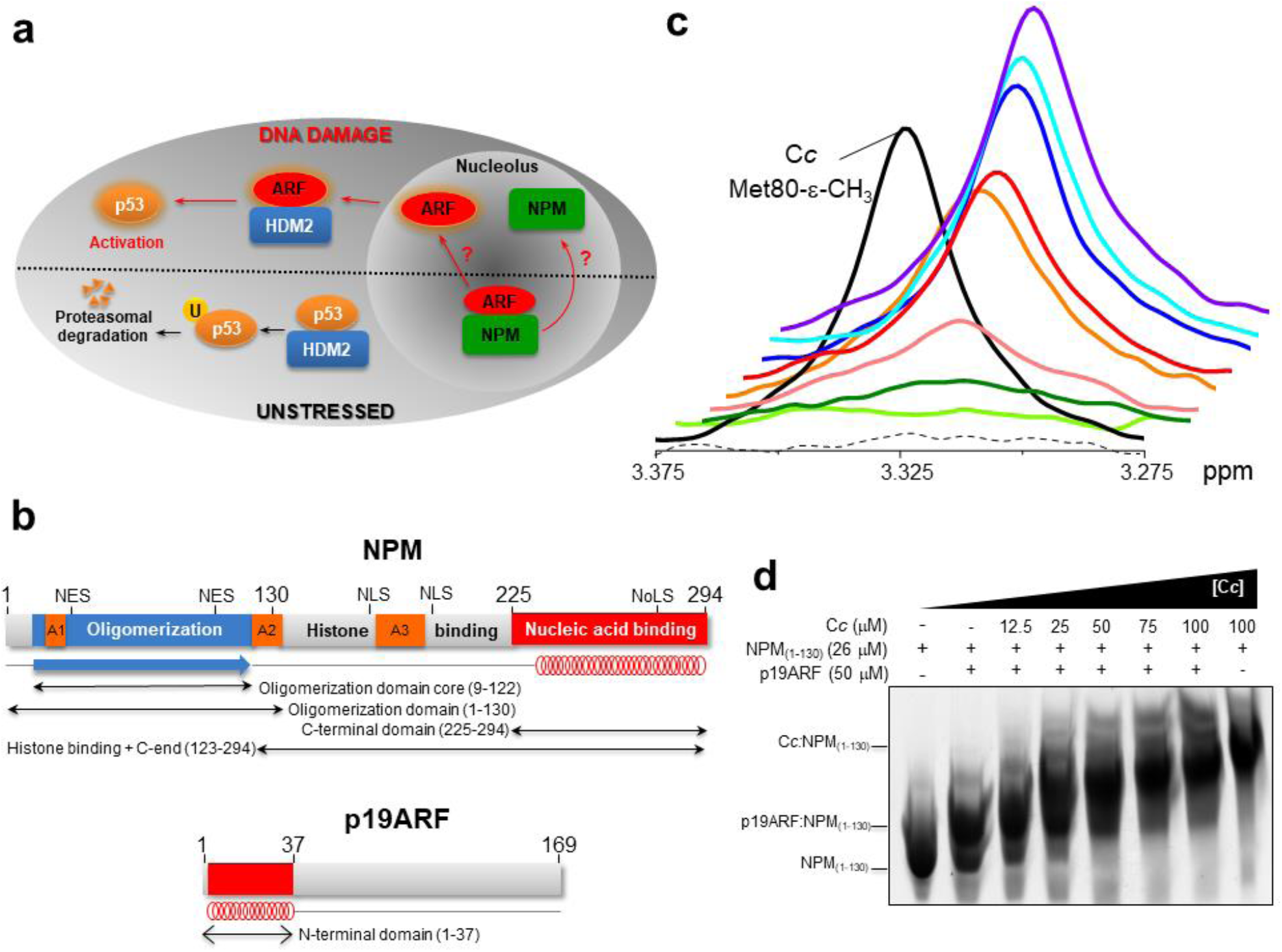
Competition between p19ARF and C*c* for NPM binding. **a**, Regulation of the ARF-p53 tumor suppressor pathway by NPM. In unstressed cells NPM blocks ARF in the nucleolus, resulting in the HDM2-mediated ubiquitination and degradation of p53. Upon DNA damage, ARF is released from the NPM-ARF complex and sequesters HDM2 in the nucleoplasm, thus preventing p53 ubiquitination. **b**, Domain organization of NPM and p19ARF. The schematic structure of NPM shows the N-terminal core domain responsible for oligomerization (blue), followed by the disordered region involved in histone binding (grey) and the C-terminal domain for nucleic acid binding (red). The acidic tracts are shown in orange. NES: nuclear export signal, NLS: nuclear localization signal, NoLS: nucleolar localization signal. The schematic structure of p19ARF shows its folded N-terminal domain. The length and secondary structure elements of the NPM constructs and the p19ARF peptide used in this work are also presented. **c**, 1D ^1^H NMR spectra monitoring Met-80 methyl signal of 13 μM reduced C*c* either free (solid black) or upon successive additions of 6 μM NPM_(1-294)_ (dotted line) and the p19ARF peptide at 41.5 μM (light green), 43 μM (dark green), 44 μM (pink), 45 μM (orange), 50 μM (red), 55 μM (blue), 60 μM (cyan) and 75 μM (violet). Spectra were shifted to show broadening of the Met80-εCH_3_ signal. **d**, EMSA showing the competitive interaction between NPM_(1-130)_ and p19ARF_(1-37)_ in the presence of C*c*. Migration of NPM_(1-130)_ either free or in complex with C*c* or p19ARF_(1-37)_ are indicated.

The ^1^H nuclear magnetic resonance (NMR) signal of the Met80-εCH_3_ of reduced C*c* senses any interaction of this protein with binding partners (*22*). Notably, this signal broadened beyond detection limits upon addition of 6 μM full-length NPM (NPM_(1-294)_) onto a 13 μM C*c* sample (Fig. 2c). Titration of the p19ARF_(1-37)_ domain — responsible for NPM binding (*23-25*) — onto the C*c*:NPM mixture gradually restored the Met80-εCH_3_ signal (Fig. 2c). Hence, the p19ARF competes with C*c* for NPM binding. Likewise, the Met80-εCH_3_ signal of C*c* disappeared upon adding the NPM oligomerization domain core (NPM _(9-122)_), known to bind p19ARF (*12*) showing both proteins, Cc and p19ARF, bind to the oligomerization domain core. Then, addition of p19ARF_(1-37)_ to the mixture partially rescued the NMR signal (Fig. S3). At high p19ARF_(1-37)_ concentrations, protein aggregation prevented further recovery.

Competition between p19ARF_(1-37)_ and C*c* for the NPM oligomerization domain (NPM_(1-130)_) was also observable in electrophoretic mobility shift assays (EMSA). Free NPM_(1-130)_ ran rapidly across the gel, but lagged in presence of p19ARF_(1-37)_ or C*c* (Fig. 2d), the latter inducing the largest shift. Interestingly, progressive additions of C*c* to the initial p19ARF_(1-37)_:NPM_(1-130)_ mixture resulted in retardation patterns indicating a change in complex composition from p19ARF_(1-37)_:NPM_(1-130)_ to C*c*:NPM_(1-130)_ (Fig. 2d). Hence, C*c* and p19ARF compete for binding to the oligomerization domain of NPM.

### NPM uses its oligomerization domain to recognize C*c* and p19ARF

Isothermal titration calorimetry (ITC)-derived dissociation constant (*K*_D_) values (Table S1 and Fig. S4a-d) indicate C*c* shows similar affinities towards the full-length protein (NPM_(1-294)_) and its oligomerization core (NPM_(9-122)_). The C*c*:NPM stoichiometry was 2:1 in both cases, with no binding cooperativity indeed. Titration analysis using a general binding model based on the overall binding constants was performed according to ref. *26*. Removal of the oligomerization domain in the NPM_(123-294)_ construct, which keeps the histone binding region (*27*), increases the *K*_D_ value to 13 μM. Further, C*c* shows low affinity (*K*_D_ *ca*. 200 μM) to the NPM_(225-294)_ construct— responsible for nucleic acid binding (*28*) and nucleolar location (*29*). Hence, C*c* preferentially binds NPM by its oligomerization domain and disordered histone binding region. When high ionic conditions were employed, a single binding site for C*c* in NPM_(1-294)_ or NPM_(9-122)_ were observed (Table S1 and Fig. S5a-b). Noteworthy, the estimated values for enthalpy and entropy did not contain any extrinsic contribution from buffer de/ionization as a buffer with negligible ionization enthalpy was employed in the ITC experiments. Those parameters can thus be considered as intrinsic thermodynamic parameters for the interaction of C*c* with its partners. Interestingly, ITC titrations using oxidized C*c* resulted in similar binding affinities as those attained when using reduced C*c* (Table S1 and Fig. S5c-d).

Notably, kinetic dissociation constant (*k*_off_) values obtained by surface plasmon resonance (SPR) indicate the half-life of C*c*:NPM_(1-294)_ complex is several fold that of C*c*:NPM_(9-122)_ (Fig. S4e). Indeed, electrostatic interactions between C*c* and the A2 region of NPM (see below) would hinder dissociation. NPM tails may affect such kinetics by steric hindrance and/or because the entropic cost of restraining their dynamics.

The interactions between NPM constructs and p19ARF_(1-37)_ yield rather complex thermograms (Fig. S4f-g), displaying three apparent phases when NPM_(1-294)_ is the analyte. Fitting them considering three classes of independent binding sites would bias the analysis of the first titration phase. Restricting the fit to this one, yielded a *K*_D_ value of 0.28 μM and a stoichiometry of 5.6 using a single class of independent binding sites (Fig. S4f, Table S1). NPM_(9-122)_ binds p19ARF_(1-37)_ with the same affinity as NPM_(1-294)_ (Fig. S4g, Table S1). Despite their different affinities towards NPM, C*c* and p19ARF_(1-37)_ may compete during an eventual accrual of C*c* in the nucleus after DNA damage (*16*). Notably, binding of the unstructured and flexible p19ARF_(1-37)_ to the NPM constructs was entropy driven, as found in many complexes involving intrinsically disordered proteins (*30-32*).

### Electrostatic interactions are key in cytochrome *c* and nucleophosmin complex formation

Because of its high pI, C*c* may interact with negative NPM regions. According to *K*_D_ values, the affinities of C*c* towards NPM_(1-294)_ and NPM_(9-122)_ decrease by 12-fold and 3-fold, respectively, upon addition of 0.1 M KCl (Fig. S5a-b, Table S1). In agreement, the affinity loss correlates with increases in binding enthalpy. The larger ionic strength effects observed with NPM_(1-294)_ may reflect electrostatic repulsion between NPM disordered regions aiding C*c* access to its acidic binding site at the oligomerization ring and the nearby disordered acidic, histone binding region of NPM (Fig. 2b). At high ionic strength, the stoichiometry of the complex formed by C*c* with both, NPM_(1-294)_ and NPM_(9-122)_, was 1:1 (Table S1). Hence, one of the binding sites observed at low ionic-strength values may correspond to unspecific electrostatic interactions.

Heteronuclear single-quantum correlation (HSQC) spectra of ^15^N labelled C*c* samples, before and after the addition of NPM_(1-294)_ (Fig. 3a) allowed us to measure averaged values (Δδ_avg_) for chemical-shift perturbations (CSP). Notably, 11 amide resonances (residues G6, K7, K8, Q16, G24, H33, G77, T78, I81, K86 and E89) shifted substantially (Δδ_avg_ > 0.05 ppm) (Fig. 4b). Others underwent smaller yet significant perturbations. Most shifted signals mapped to fringes of the heme cleft (Fig. 3b), as in other interactions with C*c* partners (*14, 33, 34*). Four lysine residues located at the interface. Addition of the N-terminal core domain of NPM (NPM_(9-122)_) to C*c* samples induced significant but smaller changes in 21 amide C*c* resonances, consistently with the higher *K*_D_ value (Fig. S6a) spread across the same surface patch. Signals either shifted or broadened (Fig. S6a-b). Again, the interaction surface includes ionic residues: five lysines (5, 7, 8, 13 and 86) and two acidic residues (E89 and E93).

**Fig. 3.**
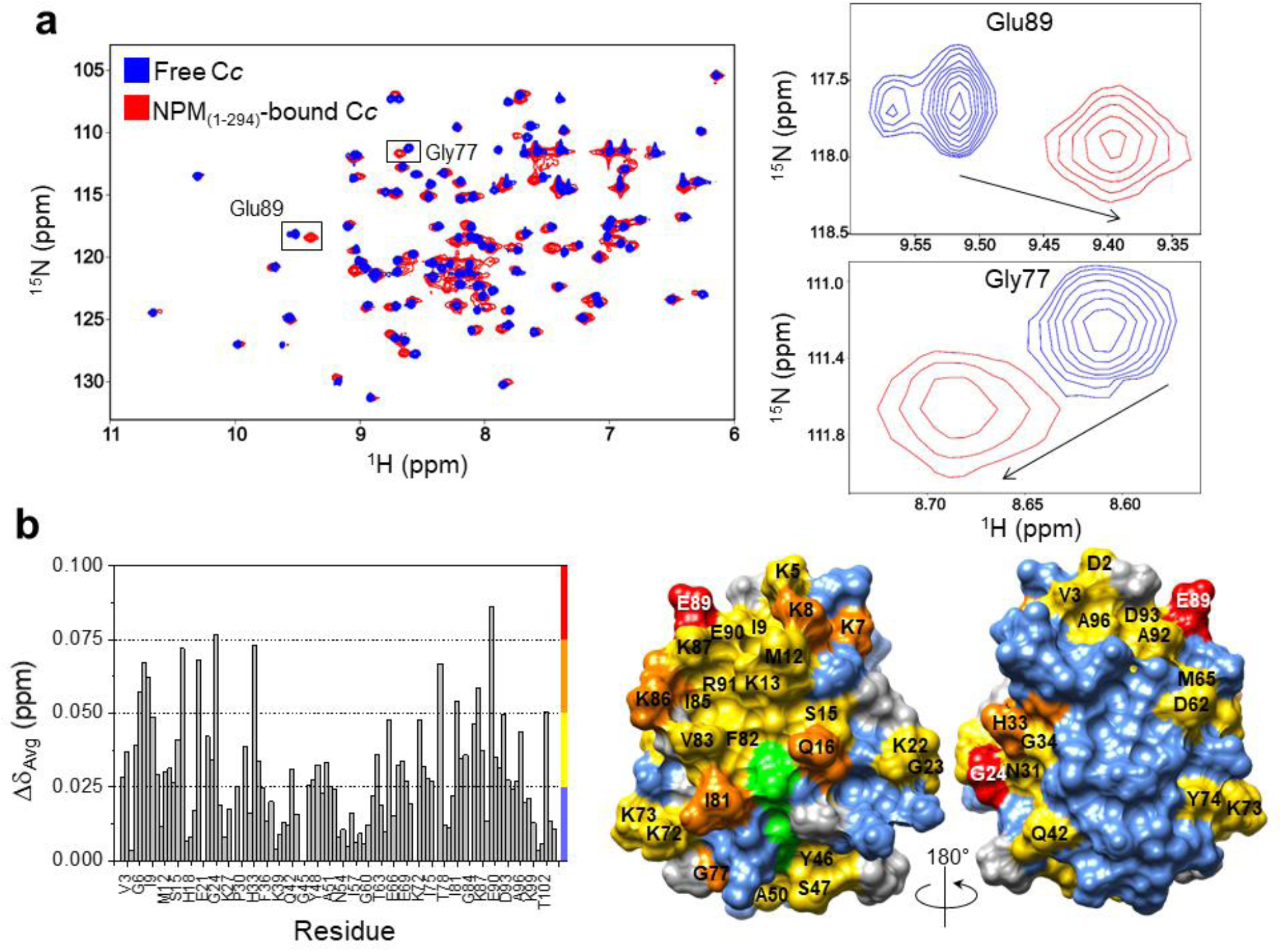
NMR titrations of ^15^N-labeled C*c* with NPM_(1-294)_. **a, *Left***, Superimposed [^1^H-^15^N] 2D HSQC spectra of C*c*, which is either free (blue) or bound to NPM_(1-294)_ (red) at the C*c*:NPM_(1-294)_ molar ratio of 1:0.5. ***Right***, Close-up views of the spectral signals for Glu89 and Gly77, arrows indicate the direction of CSPs. **b, *Left***, Average CSPs (Δδ_Avg_) of C*c* in complex with NPM_(1-294)_, at a C*c*:NPM_(1-294)_ ratio of 1:0.5. Color bars represent Δδ_Avg_: insignificant (< 0.025 ppm [blue]), small (0.025-0.050 ppm [yellow]), medium (0.050-0.075 ppm [orange]) and large (>0.075 ppm [red]). ***Right***, Mapping of C*c* residues perturbed upon binding with NPM_(1-294)_. C*c* is rotated 180° around vertical axes. Residues are colored according to their Δδ_Avg_ values (ppm), with the same-colour code. Non-assigned residues and prolines are in grey. The heme group is in green.

**Fig. 4.**
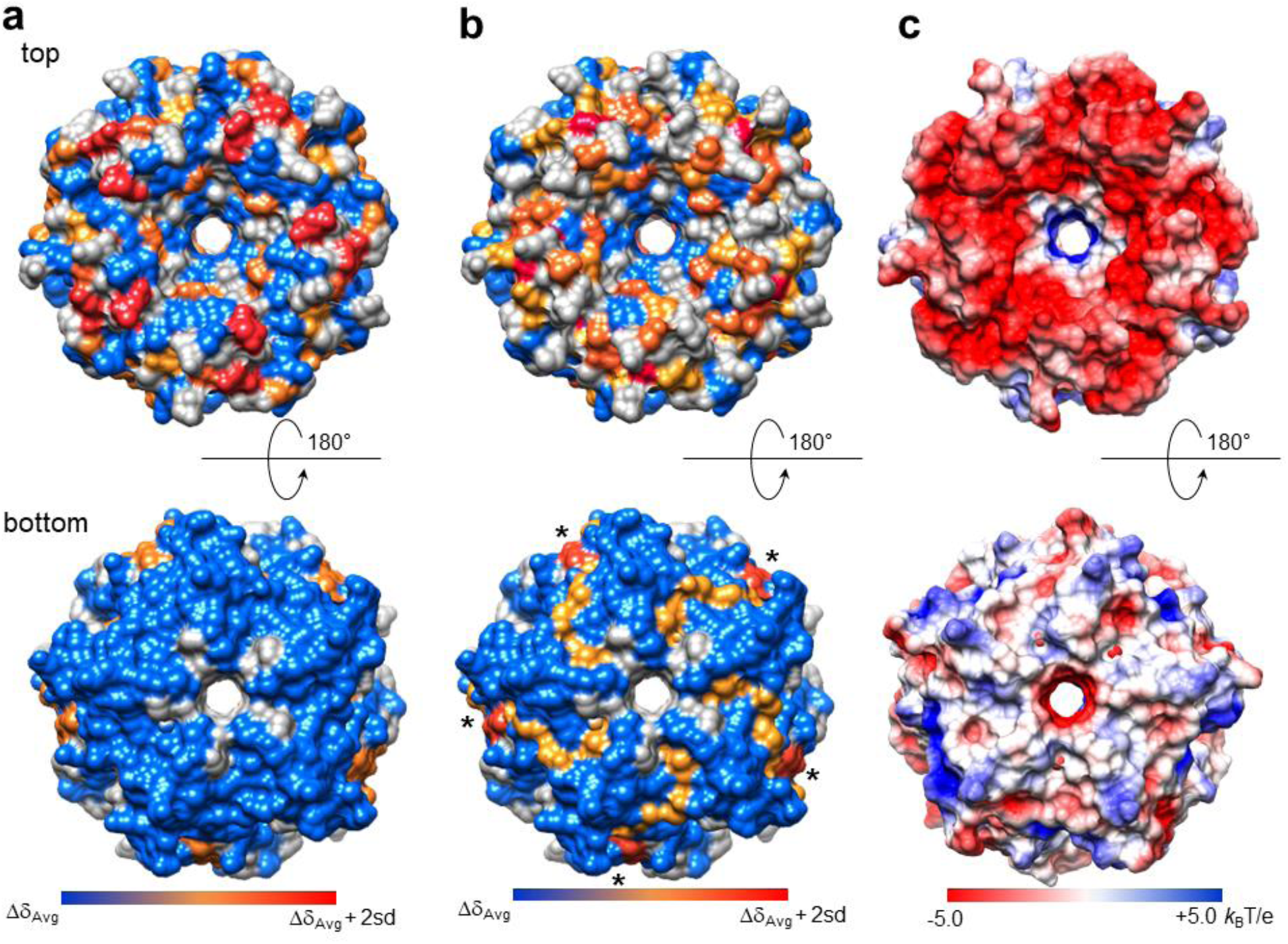
NMR titrations of ^2^H-^15^N-labeled NPM_(1-130)_ with C*c* or p19ARF_(1-37)_. **a**,**b**, Mapping of NPM_(1-130)_ surface residues perturbed upon binding to C*c* (**a**) or p19ARF_(1-37)_ (**b**). Molecules are rotated 180° around vertical axes in each view. Residues are colored according to their averaged CSPs (Δδ_Avg_): insignificant (Δδ < [Δδ_Avg_ + Sn-1], blue), medium ([Δδ_Avg_ + Sn-1] ≤ Δδ < [Δδ_Avg_ + 2Sn-1], orange) and large (Δδ > [Δδ_Avg_ + 2Sn-1], red). Non-assigned residues and prolines are in grey. Asterisks indicate interfaces between monomers of NPM. **c**, The electrostatic potential map of NPM_(1-130)_ was calculated in Chimera with Adaptive Poisson-Bolzmann Solver (APBS) at 0.1 M ionic strength.

Conversely, we recorded TROSY (transverse relaxation optimized spectroscopy) spectra of a ^2^H-^15^N-labeled sample of NPM_(1-130)_. Some NPM resonances shifted upon C*c* addition at a molar C*c*:NPM_(1-130)_ ratio of 2:1 (Fig. S7a). Most residues showing significant CSPs clustered at the acidic “top patch” of the oligomerization domain (Fig. 4a,c). Likewise, p19ARF_(1-37)_ protein binds the same region of NPM_(1-130)_ (Fig. 4b), despite CSPs being smaller (Fig. S7b-d). Nevertheless, unlike C*c*, p19ARF_(1-37)_ binds the interfaces between the NPM monomers, also interacting with residues at the opposite face of NPM_(1-130)_ (Fig. 4b, indicated by asterisks). This fuzziness, in line with our ITC data, may explain NPM_(1-130)_ undergoing smaller CSPs in the presence of p19ARF_(1-37)_ (Fig. S7c-d). The eventual impact of His-tag on the core structure of NPM can be discarded since no significant structural changes were observed (root-mean-square deviation of backbone atoms is 0.35 Å; Fig. S8) when the two pentameric NPM X-ray structures available at the Protein Data Bank—with (2P1B, *35*) and without His-tag (4N8M, *25*)—are compared. We thus corroborated that His-tagged NPM_(1-130)_ is properly folded as inferred from NMR and CD (circular dichroism) experiments (Fig. S9a,b). These findings agree with the SDS-PAGE assays performed with His-tagged NPM_(1-130)_ samples, which show an intense majority band corresponding to the pentameric form at any ionic strength (Fig. S9c).

### Three-dimensional structures of NPM and the C*c*:NPM complex

We resorted to X-ray diffraction to get further structural data of the C*c*:NPM complex. C*c* and NPM_(9-122) —_the oligomerization domain core— were co-crystallized using the microbatch technique in 25% PEG 400, 0.1 M MES, 0.1 M MgCl_2_ and 0.1 M TCEP as an additive. The refined model (PDB 5EHD) contained 20 monomers of NPM_(9-122)_ in the asymmetric unit, arranged in four pentamers as previously observed (*35*). Residues 9-14, 119-122 and the C-terminal His-tag were invisible in the electron density map. Each NPM_(9-122)_ monomer forms a compact domain comprising an eight-stranded β-barrel of jellyroll topology (Fig. 5a), as previously found (*35*) (PDB 2P1B). No electron density associated with C*c* was noticeable in any of the four NPM pentamers, despite its presence in the co-crystal being supported by SDS-PAGE (Fig. S10).

**Fig. 5.**
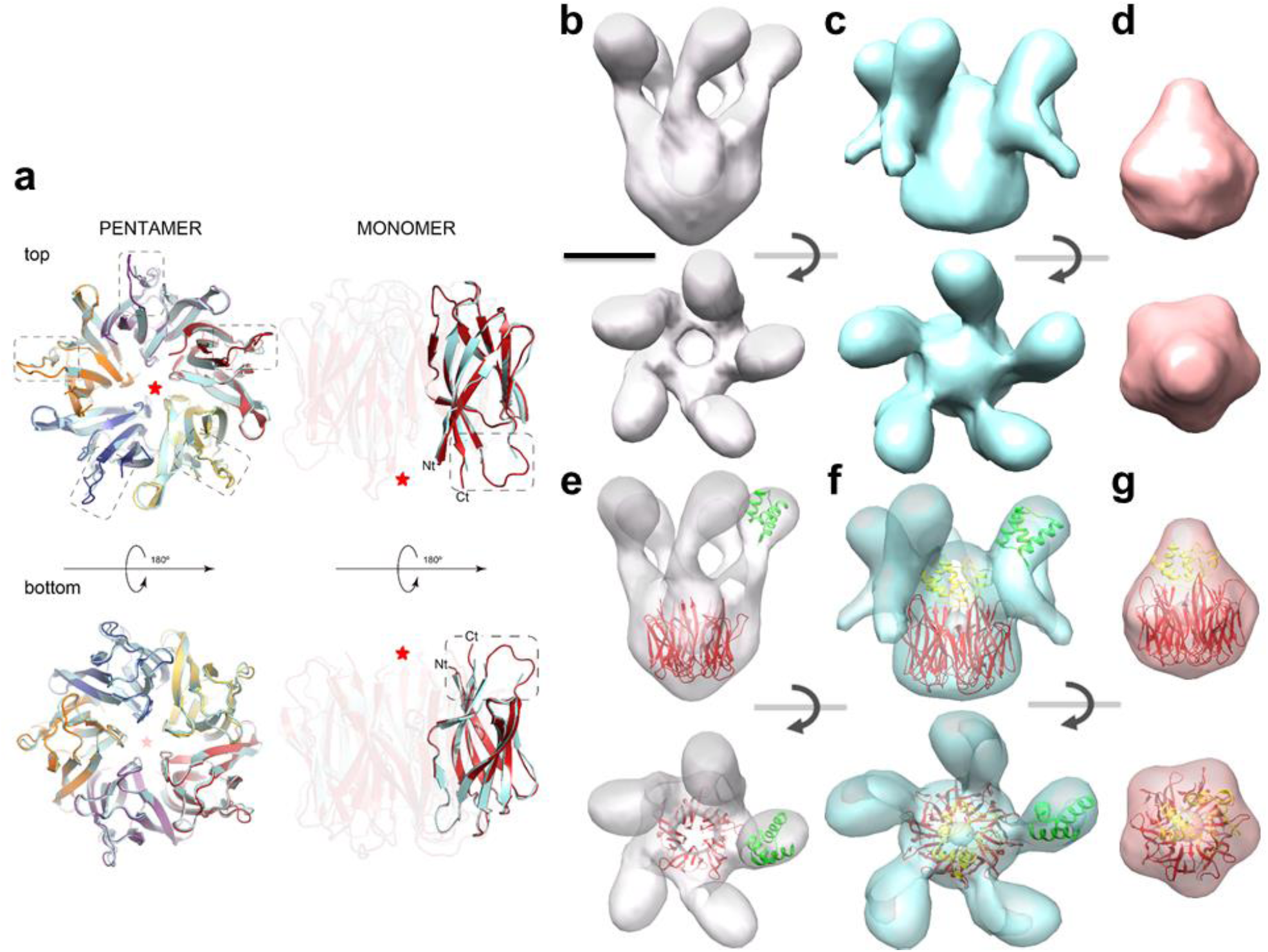
X-ray and EM 3D structures of NPM_(9-122)_ and NPM_(1-294)_ in complex with C*c. Left*, Superimposition of the pentameric crystal structures of NPM_(9-122)_, either free (PDB 2P1B) (*35*) or bound to C*c* (this work). The monomers of free NPM_(9-122)_ are all in light blue, whereas those of the C*c:*NPM_(9-122)_ complex are in different colors. The dotted squares represent ordered loops traced due to the presence of the C*c*. The red asterisk indicates the C*c* binding site. ***Right***, Three-dimensional reconstruction of NPM_(1-294)_ and the C*c*:NPM_(1-294)_ and C*c*:NPM_(9-122)_ complexes by EM. **a-c**, Side (*top*) and end-on (*bottom*) views of the 3D reconstructions of NPM_(1-294)_ (**a**), C*c*:NPM_(1-294)_ (**b**) and C*c*:NPM_(9-122)_ (**c**). **d-f**, The same views after docking the crystal structures of NPM_(9-122)_ (red, this work); the globular, C-terminal domain of NPM (NPM_(225-294)_) (green, PDB 2VXD), and C*c* (yellow, PDB 1J3S). Scale bar is 5 nm.

One face of the NPM_(9-122)_ pentamer contains the N- and C-termini (top view of Fig. 5a), and the flexible loops connecting β2 and β3 (highlighted by dotted lines) comprising the residues D34 to E39 of the acidic tract A1 (Fig. 2b). These loops face C*c* in our NMR analysis (Fig. 4a) and are evident in the C*c*:NPM_(9-122)_ electron density map whereas being invisible in the published structure of free NPM_(9-122)_ (*35*). The lack of C*c* electron density in the maps suggests this protein assumes multiple orientations across the crystal.

We then resorted to electron microscopy (EM) to get structural information on NPM_(1-294)_, as well as the complexes C*c*:NPM_(1-294)_ and C*c*:NPM_(9-122)_. Because of their small size and flexibility, negative staining was used. This technique, because of its low-resolution, cannot describe the nature of the interaction between NPM and C*c* but has nevertheless provided us with clear information on where C*c* binds. The images of NPM_(1-294)_ (162 kDa) revealed a major, end view often showing a clear five-fold symmetry (Fig. S11a) and a scant one revealing a base structure from which several, thin and flexible masses protrude. 28637 particles were used for 3D reconstruction. C5 symmetry was applied since NPM is a homopentamer. The volume obtained showed (Fig. 5b) a base of ∼5 nm width and 4.5 nm height from which five arms of 5 nm length protrude, with a ∼3 nm globular mass at the tip of each arm. Our crystal structure of NPM_(9-122)_ fits well into this 3D reconstruction. Likewise, the atomic structure of the last 51 residues of NPM (*36*) (PDB 2VXD, residues 243-294) docks well into the globular arm-tip domains (Fig. 5e). The disordered middle region of NPM (residues 123-242) links the two structural domains (Fig. 2b). As expected for an EM reconstruction, the NPM His-tag cannot be visualized due to low resolution and its high mobility.

When staining a C*c*:NPM_(1-294)_ mixture, EM views were very similar to apo-NPM_(1-294)_ ones, displaying the two orientations above (Fig. S11b). Outwardly, no obvious differences marked the C*c* molecule bound to the NPM_(1-294)_ pentamer, indicating the C*c* molecule lodges within the cavity formed by the NPM_(1-294)_ arms. The 3D reconstruction of the complex, given its small size (174 kDa) and flexibility, required 29500 particles. We imposed C5 symmetry throughout the 3D reconstruction procedure despite the inaccuracy for reconstructing a single C*c* molecule in the complex. Nevertheless NPM_(9-122)_ and NPM_(1-294)_ variants bind C*c* with similar affinities (Table 1), suggesting the N-terminal core domain of NPM is the principal C*c* interaction site and has five identical, potential binding sites.

The resulting volume (Fig. 5c) resembled that observed for apo-NPM_(1-294)_, displaying a central mass and five protruding arms ending in small globular masses. Noteworthy, there are two important differences between the C*c*:NPM_(1-294)_ complex and free NPM_(1-294)_: first, a retraction in the arms (∼2 nm) closer to the core mass and second, a protrusion in the core partially filling the central cavity and touching the base of the arms. This bulge discloses the C*c* molecule attached to the central cavity, in contact with the NPM core domain, and induces a contraction of the arms by interacting with their base. Such a contraction seems to induce the lateral protrusion of part of the flexible middle region of the chaperone. Only a single C*c* molecule is bound to the NPM_(1-294)_ pentamer in this EM structure (Fig. 5c,f) obtained at high ionic strength — ca. 150 mM before drying. Indeed, ITC data indicate a 1:1 stoichiometry for the interaction between C*c* and the NPM_(1-294)_ oligomer at high ionic strength (Table S1).

The atomic structures of the NPM_(9-122)_ pentamer and the C-terminal globular domain dock well into the corresponding positions of the volume (Fig. 5f). NMR-restrained docking calculations enabled us to fit the C*c* molecule in the EM protruding mass. CSPs obtained from NMR analysis of the NPM_(9-122),_ complex were used as input data. High-score models placed C*c* towards the highly acidic face of NPM_(9-122)_, as shown for the best-scoring solution (Fig. S12). In this one, the heme cleft points to the acidic stretches of NPM_(9-122)_. Docking results were consistent with C*c* lodging in the EM protrusion contacting the inner side of the NPM core domain.

Since NPM_(9-122)_ binds C*c* with the same stoichiometry as NPM_(1-294)_ (Table S1), we sought to confirm the location of C*c* in the complex. Images of negatively stained C*c*:NPM_(9-122)_ complex (Fig. 5d) revealed a small, globular structure consistent with the molecular mass of the complex (84 kDa). 21215 particles were used for the 3D reconstruction, and C5 symmetry was again imposed. The volume obtained (Fig. 5d) displayed the base alike the chaperone core NPM_(1-294)_ and the C*c*:NPM_(1-294)_ complex. Resembling the C*c*:NPM_(1-294)_ complex, a small mass protrudes from the base of the pentamer, which can be clearly attributed to bound C*c*. This reading invigorates once NPM_(9-122)_ and C*c* dock in their corresponding places (Fig. 5g).

All these findings point to C*c* binding with NPM_(1-294)_ through the internal part of the NPM core (NPM_(9-122_), assisted by some regions at the base of the inner side of the chaperone arms, probably inducing their contraction. These results also explain why, even though the NPM pentamer contains five identical binding sites for C*c*, only one molecule binds to the chaperone: once the first C*c* molecule binds to one of the five potential binding sites, there is no room for the others.

### NPM undergoes phase separation with ARF or C*c*

NPM is known to undergo liquid-liquid phase separation via heterotypic interactions with nucleolar components (e.g., proteins displaying arginine-rich (R-rich) motifs) able to interact with the NPM acidic tracts (*37,38*). Since ARF – which contains R-rich motifs – interacts with NPM, we addressed its ability to promote phase separation of NPM *in vitro*. Titration of NPM_(1-294)_ with p19ARF_(1-37)_ resulted in formation of phase separated droplets (Fig. 6a, *left*), as does NPM with human p14ARF to form gel-like condensates (*39*). This process was observed when the p19ARF_(1-37)_:NPM_(1-294)_ ratio increased up to 5:1, being clearly evident at 10:1 (Fig. 6a, *left*). NPM_(1-294)_ phase separated with p19ARF_(1-37)_ or C*c* also in the presence of 100 mM KCl (Fig. S13). Phase separation occurs after significant saturation in the p19ARF_(1-37)_:NPM_(1-294)_ complex. This could explain the rather complex thermograms showed by NPM in complex with p19ARF_(1-37)_, as discussed above (Fig. S4f-g). We likewise observed phase separation when using Oregon Green 488-labeled NPM_(1-294)_ and p19ARF_(1-37)_ labeled with rhodamine (Fig. 6b, *upper*). Remarkably, C*c* does not contain R-rich motifs but likewise mediated phase separation of NPM_(1-294)_ starting at the 5:1 C*c*:NPM_(1-294)_ ratio (Fig. 6a, *right*), and labeled NPM_(1-294)_ showed phase separation in complex with Alexa Fluor 647-labeled C*c*, which exhibits signs of spinodal decomposition (Fig. 6b, *lower*). The fact that ARF and NPM form heterotypic phase separated droplets is in agreement with ARF sequestration by NPM in the nucleolus (*3*), whereas the herein reported finding that C*c* is able to form phase separated droplets with NPM is consistent with the novel interaction detected between the hemeprotein and NPM in the nucleoli (Fig. 1c).

**Fig. 6.**
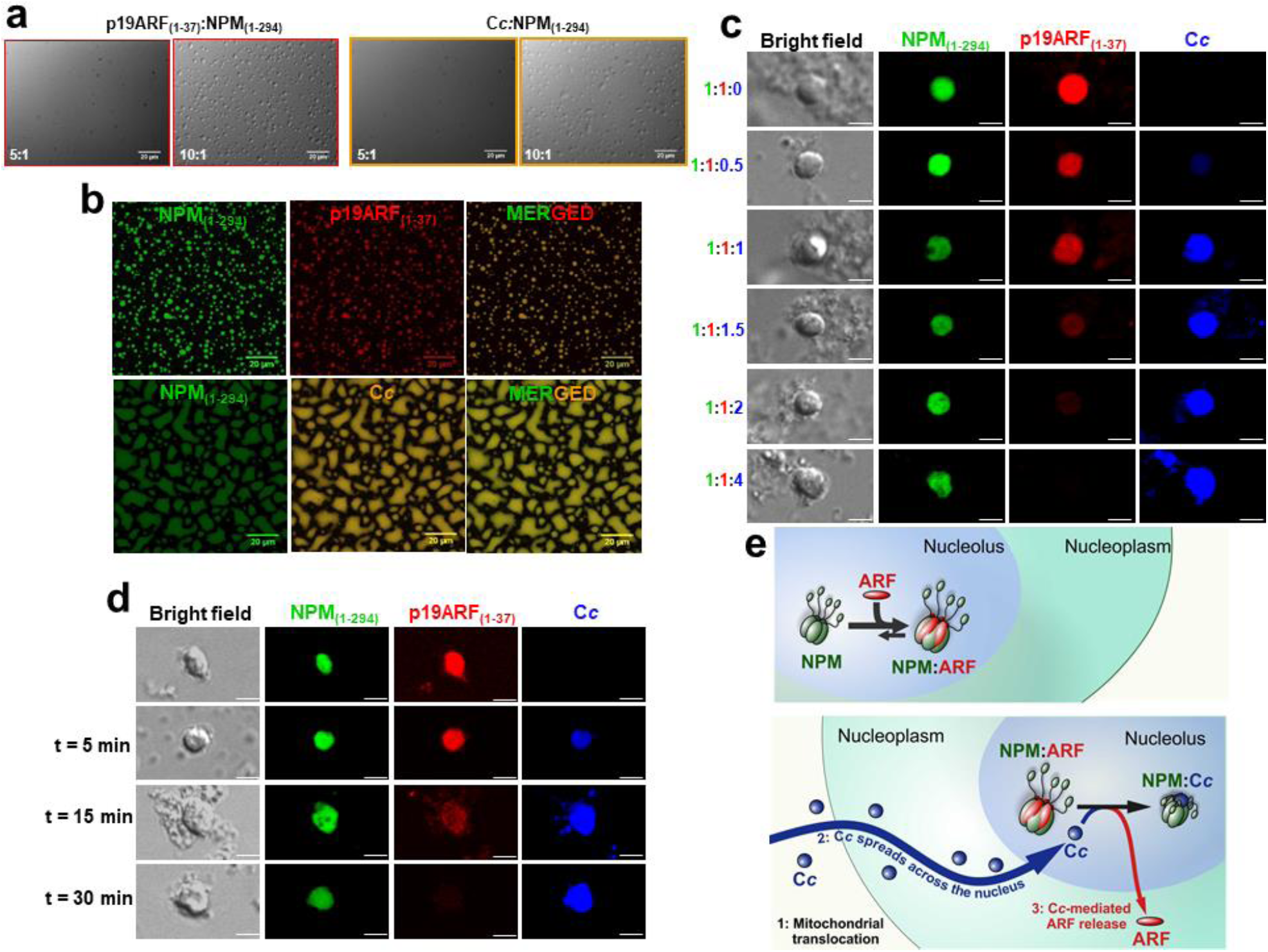
Liquid-like properties of NPM in complex with either p19ARF or C*c*. **a**, *In vitro* droplets formed by NPM_(1-294)_ with either p19ARF_(1-37)_ (*left*) or C*c* (*right*) showed by DIC microscopy images. NPM_(1-294)_ was at 5 μM, whereas p19ARF_(1-37)_ and C*c* were added at p19ARF_(1-37)_:NPM_(1-294)_ and C*c*:NPM_(1-294)_ ratios of 5:1 and 10:1, respectively. **b**, Fluorescence confocal microscopy images of phase separation by 5 μM NPM_(1-294)_ with either 75 μM p19ARF_(1-37)_ (*upper*) or 75 μM C*c* (*lower*). NPM_(1-294)_ was labeled with Oregon Green 488 (green channel), whereas p19ARF_(1-37)_ with NHS-Rhodamine (red channel) and C*c* with Alexa Fluor 647 (far-red channel, depicted in orange). Droplets were imaged in 10 mM Tris buffer (pH 7.5) containing 100 mM NaCl and 2 mM DTT. **c**, Confocal microscopy images of isolated nucleoli incubated with 5 μM Orange Green-NPM_(1-294)_ (green channel) and 5 μM Rhodamine-p19ARF_(1-37)_ (red channel), before and after titration with increasing concentrations (ranging from 2.5 to 20 μM) of Alexa Fluor 647-C*c* (far-red channel, depicted in blue). **d**, Confocal microscopy images of isolated nucleoli pre-incubated with 5 μM NPM_(1-294)_ and 5 μM p19ARF_(1-37)_ upon addition of 5 μM C*c* over time. Scale bars are 3 μM. All images were taken with a 63x objective (oil). **e**, Proposed molecular mechanism of C*c*-induced release of the ARF tumor suppressor from NPM. ***Upper***, under homeostasis, NPM and ARF form liquid condensates, keeping ARF sequestered in the nucleolus. ***Lower***, upon DNA damage, C*c* enters the nucleolus where it competes with ARF for NPM binding, releasing ARF from the nucleolus.

### C*c* liberates ARF from the nucleolus

To test the relevance of such *in vitro* findings under physiological conditions, we monitored how Alexa Fluor 647-C*c* could interfere with Oregon Green 488-NPM_(1-294)_ and Rhodamine-p19ARF_(1-37)_ accumulation in isolated mammalian nucleoli. As observed in Fig. 6c (*upper*), both NPM and p19ARF_(1-37)_ are incorporated within nucleoli in the absence of C*c*. However, in the presence of increasing concentrations of Alexa Fluor 647-C*c*, the fluorescence of p19ARF_(1-37)_ gradually diminished up to disappear while that of NPM_(1-294)_ remains unalterable (Fig. 6c). Addition of C*c* to pre-incubated p19ARF:NPM nucleoli also resulted in a decrease in the p19ARF_(1-37)_ fluorescence over time (Fig. 6d). These results clearly indicate that p19ARF is in fact released from the nucleoli when C*c* is incorporated to the membrane-less organelle (Fig. 6e).

## Discussion

Nucleoli are key organelles in the cell stress response and different stimuli cause the loss of function of such essential membrane-less structures (*40,41*). A crucial element in this process is NPM, which is responsive to stress of different kinds (*42, 43*). NPM regulates the ARF/p53 pathway by keeping ARF sequestered in the nucleoli but releasing it upon DNA damage (*12*). We herein demonstrate that endogenous C*c* is translocated into the nucleus in both cancer and non-cancer cells, thereby interacting with endogenous NPM, upon DNA insults.

At physiological ionic strength, a single C*c* molecule binds to the oligomerization domain of NPM. The middle-disordered region of NPM provides a kinetic barrier for the interaction with C*c*, without affecting the affinity. Our crystal (C*c*:NPM_(9-122)_) and EM structures (C*c*:NPM_(9-122)_ and C*c*:NPM_(1-294)_) disclose that binding of C*c* to the NPM oligomerization core strongly affects its loops and the intrinsically disordered middle region of full-length NPM. This region, essential for histone chaperone activity and heterotypic transitions of NPM (*37*), shrinks and approaches to the core domain to which C*c* is bound. The 3D structure herein displays C*c* hinders the access to the acidic tracts of the disordered binding region of NPM, to which several partners bind (*38*). Our biophysical data clearly indicate that C*c* can oust p19ARF_(1-37)_ from NPM. The CSPs maps of NPM in the C*c*:NPM and ARF:NPM complexes visibly display an overlap between their respective binding sites on NPM.

NPM is a well-known nucleolar marker, which is able to undergo liquid-liquid phase separation (LLPS) by homo- and heterotypic interactions and greatly contributes to the liquid-like features of nucleolus (*37,38*). Recent data has highlighted the key role of linear R-rich elements in the proteins exhibiting heterotypic interactions with NPM (*37*), yet K-rich stretches can also be involved in the assembly of other membrane-less organelles (e.g. stress granules) (*44*). According to Kriwacki and co-workers (*37*), heterotypic interactions require extended NPM conformations, which do occur at physiological ionic strength. Such interactions are crucial to enable recognition by RNA and assembly of pre-ribosomal particles (*37*). On the other hand, homotypic interactions have been detected under non-physiological, low-salt conditions, in which a compact conformation of NPM is populated. The balance between both types of interactions is proposed to be essential for keeping the “liquid-like scaffold of the granular component of nucleoli” (*37*). Our data herein provides a mechanism by which a small, K-rich globular protein such as C*c* triggers the transition from an extended state to a compact conformation of NPM under physiological, high salt conditions. In this regard, it is worth noting the differences in shape and physical features between NPM droplets induced by ARF and C*c* (Fig. 6a,b). We thus describe C*c* as a cofactor modulating the conformational equilibrium of NPM, along with its multi-valence and LLPS mechanism. C*c-*induced homotypic condensation of NPM could be a reversible switch for regulation of ribosomal biosynthesis under DNA damage and release of tumor suppressor factors like ARF.

In agreement with the NPM ability to form droplets not only with ARF but also with C*c*, the hemeprotein is incorporated into the nucleoli so as to shift ARF out from the preformed NPM-ARF ensembles and from the nucleoli themselves, thereby explaining the molecular mechanism underlying the ARF release and its later activation upon DNA damage.

On the basis of our data, we propose that the C*c*:NPM nucleolar interaction, involving the extended-to-compact transition of NPM and release of NPM-sequestered ARF, can be expanded to a general mechanism for DNA damage response in which the K-rich regions of C*c* act in a pleiotropic, moonlighting manner on histone chaperones.

## Acknowledgements

The authors thank Dr. J. Martínez-Fábregas (Scotland) for kind help with proteomic analyses. **Funding:** The authors are grateful to the Spanish Ministry of Economy and Competitiveness (BIO2015-70092-R, BFU2015-71017/BMC, BFU2017-90030-P, BFU2016-75984/BMC, PGC2018-096049-B), the European Commission (European Regional Development Fund and European Research Council (CONCERT, contract number 648201), the Regional Government of Andalusia (BIO198), the Ramon Areces Foundation, the NMR Facility at CITIUS (University of Seville) and Waters-TA Instruments. This work has been supported by iNEXT, grant number PID 3407, funded by the Horizon 2020 programme of the European Commission. C.A.E.R., M.A.C.C., A.V.-Cruz and E.S thank Cámara Foundation, the Spanish Ministry of Education, Culture and Sports (FPU18/06577 and FPU16/01513) and La Caixa foundation for their respective PhD fellowships. IRB Barcelona is recipient of a Severo Ochoa Award of Excellence from MINECO (Government of Spain).

## Author contributions

K.G.A., A.D.Q., I.D.M. and M.A.R. designed the research; K.G.A., M.A.C.C. carried out the proteomics and cellular experiments. K.G.A., C.A.E.R. and A.V.-Cruz cloned, expressed and purified proteins. K.G.A., A.D.Q. and A.V.-Campoy performed and analyzed ITC measurements. K.G.A., S.G.C., I.A. and I.D.M. performed and analyzed NMR data; K.G.A. run docking calculations. N.B.G. and J.A.H. acquired and analyzed the crystallographic data. E.S., X.S., A.D.Q. and K.G.A. implemented the phase separation assays. R.A and J.M.V. carried out the electron microscopy and the single-particle 3D reconstruction. K.G.A., A.D.Q., I.D.M. and M.A.R. wrote the paper.

## Supplementary Materials

Materials and Methods

Figures S1 to S13

Tables S1 and S2

## References

1. A. Matheu et al., Delayed ageing through damage protection by the Arf/p53 pathway. Nature 448, 375–379 (2007).

2. C. J. Sherr, Divorcing ARF and p53: An unsettled case. Nat. Rev. Cancer 6, 663–673 (2006).

3. C. Korgaonkar et al., Nucleophosmin (B23) Targets ARF to nucleoli and inhibits its function. Mol. Cell. Biol. 25, 1258–1271 (2005).

4. Y. Zhang, Y. Xiong, W. G. Yarbrough, ARF promotes MDM2 degradation and stabilizes p53: ARF-INK4a locus deletion impairs both the Rb and p53 tumor suppression pathways. Cell 92, 725–734 (1998).

5. A. Di Matteo et al., Molecules that target nucleophosmin for cancer treatment: an update. Oncotarget 7, 44821–44840 (2016).

6. Y. Yu et al., Nucleophosmin is essential for ribosomal protein L5 nuclear export. Mol. Cell. Biol. 26, 3798–3809 (2006).

7. S. Grisendi, C. Mecucci, B. Falini, P. P. Pandolfi, Nucleophosmin and cancer. Nat. Rev. Cancer 6, 493–505 (2006).

8. V. Swaminathan, A. H. Kishore, K. K. Febitha, T. K. Kundu, Human histone chaperone nucleophosmin enhances acetylation-dependent chromatin transcription. Mol. Cell. Biol. 25, 7534–7545 (2005).

9. K. Hanashiro, M. Brancaccio, K. Fukasawa, Activated ROCK II by-passes the requirement of the CDK2 activity for centrosome duplication and amplification. Oncogene 30, 2188–2197 (2011).

10. J. K. Box, et al., Nucleophosmin: from structure and function to disease development. BMC Mol. Biol. 17, 1–12 (2016).

11. L. Latonen, M. Laiho, Cellular UV damage responses—Functions of tumor suppressor p53. Biochim. Biophys. Acta – Rev. Cancer 1755, 71–89 (2005).

12. T. Enomoto, M. S. Lindstrom, A. Jin, H. Ke, Y. Zhang, Essential role of the B23/NPM core domain in regulating ARF binding and B23 stability. J. Biol. Chem. 281, 18463–18472 (2006).

13. F. -X. Qin et al., Knockdown of NPM1 by RNA interference inhibits cells proliferation and induces apoptosis in leukemic cell line. Int. J. Med. Sci. 8, 287–294 (2011).

14. J. Martínez-Fábregas et al., Structural and functional analysis of novel human cytochrome *c* targets in apoptosis. Mol. Cell. Proteomics 13, 1439–1456 (2014).

15. J. Martínez-Fabregas, I. Díaz-Moreno, K. González-Arzola, A. Díaz-Quintana, M. A. De la Rosa, A common signalosome for programmed cell death in humans and plants. Cell Death Dis. 5, e1314 (2014).

16. K. González-Arzola et al., Structural basis for inhibition of the histone chaperone activity of SET/TAF-Iβ by cytochrome c. Proc. Natl. Acad. Sci. U.S.A. 112, 9908–9913 (2015).

17. J. Martínez-Fábregas et al., New *Arabidopsis thaliana* cytochrome *c* partners: A look into the elusive role of cytochrome *c* in programmed cell death in plants. Mol. Cell. Proteomics 12, 3666–3676 (2013).

18. K. González-Arzola et al. Histone chaperone activity of *Arabidopsis thaliana* NRP1 is blocked by cytochrome *c*. Nucleic Acids Res. 45, 2150–2165 (2017).

19. J. C. Goldstein, N. J. Waterhouse, P. Juin, G. I. Evan, D. R. Green, The coordinate release of cytochrome *c* during apoptosis is rapid, complete and kinetically invariant. Nat. Cell. Biol. 2, 156–162 (2000).

20. S. Jacob, C. Miquel, A. Sarasin, F. Praz, Effects of camptothecin on double-strand break repair by non-homologous end-joining in DNA mismatch repair-deficient human colorectal cancer cell lines. Nucleic Acids Res. 33, 106–113 (2005).

21. T. K. Kerppola, Design and implementation of bimolecular fluorescence complementation (BiFC) assays for the visualization of protein interactions in living cells. Nat. Protoc. 1, 1278–1286 (2006).

22. 22. B. Moreno-Beltrán et al. Cytochrome *c*1 exhibits two binding sites for cytochrome *c* in plants. BBA-Bioenergetics. 1837, 1717–1729 (2014).

23. D. Bertwistle, M. Sugimoto, C. J. Sherr, Physical and functional interactions of the Arf tumor suppressor protein with nucleophosmin/B23. Mol. Cell. Biol. 24, 985–996 (2004).

24. K. Itahana et al. Tumor suppressor ARF degrades B23, a nucleolar protein involved in ribosome biogenesis and cell proliferation. Mol. Cell 12, 1151–1164 (2003).

25. D. M. Mitrea et al. Structural polymorphism in the N-terminal oligomerization domain of NPM1. Proc. Natl. Acad. Sci. U.S.A. 111, 4466–4471 (2014).

26. E. Freire, A. Schön, A. Velazquez-Campoy, in Methods in Enzymology. (Academic Press, 2009), vol. 455, pp. 127–155.

27. S. S. Gadad et al., The multifunctional protein nucleophosmin (NPM1) is a human linker histone H1 chaperone. Biochemistry 50, 2780–2789 (2011).

28. K. Hingorani, A. Szebeni, M. O. Olson, Mapping the functional domains of nucleolar protein B23. J. Biol. Chem. 275, 24451–24457 (2000).

29. J. W. Choi et al., Lysine 263 residue of NPM/B23 is essential for regulating ATP binding and B23 stability. FEBS Lett. 582, 1073–1080 (2008).

30. D. Parker et al., Role of secondary structure in discrimination between constitutive and inducible activators. Mol. Cell. Biol. 19, 5601–5607 (1999).

31. O. Abian, J. L. Neira, A. Velazquez-Campoy, Thermodynamics of zinc binding to hepatitis C virus NS3 protease: A folding by binding event. Proteins 77, 624–636 (2009).

32. H. Fraga, E. Papaleo, S. Vega, A. Velazquez-Campoy, S. Ventura, Zinc induced folding is essential for TIM15 activity as an mtHsp70 chaperone. BBA-Gen. Subjects 1830, 2139–2149 (2013).

33. B. Moreno-Beltrán et al., Structural basis of mitochondrial dysfunction in response to cytochrome *c* phosphorylation at tyrosine 48. Proc. Natl. Acad. Sci. U.S.A. 114, E3041–E3050 (2017).

34. I. Díaz-Moreno, J.M. García-Heredia, A. Díaz-Quintana, M. A. De la Rosa, Cytochrome *c* signalosome in mitochondria. Eur. Biophys. J. 40, 1301–1315 (2011).

35. H. H. Lee et al., Crystal structure of human nucleophosmin-core reveals plasticity of the pentamer-pentamer interface. Proteins: Struct., Funct., Bioinf. 69, 672–678 (2007).

36. C. G. Grummitt, F. M. Townsley, C. M. Johnson, A. J. Warren, M. Bycroft, Structural consequences of nucleophosmin mutations in acute myeloid leukemia. J. Biol. Chem. 283, 23326–23332 (2008).

37. D. M. Mitrea et al., Self-interaction of NPM1 modulates multiple mechanisms of liquid– liquid phase separation. Nat. Commun. 9, 1–13 (2018).

38. D. M. Mitrea et al., Nucleophosmin integrates within the nucleolus via multi-modal interactions with proteins displaying R-rich linear motifs and rRNA. eLife 5, e13571 (2016).

39. E. Gibbs, B. Perrone, A. Hassan, R. Kümmerle, R. Kriwacki, NPM1 exhibits structural and dynamic heterogeneity upon phase separation with the p14ARF tumor suppressor. J Magn. Reson. 310, 106646 (2020).

40. C. Lee, B. A. Smith, K. Bandyopadhyay, R. A. Gjerset, DNA damage disrupts the p14ARF-B23(nucleophosmin) interaction and triggers a transient subnuclear redistribution of p14ARF. Cancer Res. 65, 9834–9842 (2005).

41. C. P. Rubbi, J. Milner, Disruption of the nucleolus mediates stabilization of p53 in response to DNA damage and other stresses. EMBO J. 22, 6068–6077 (2003).

42. Z. Yao et al., B23 acts as a nucleolar stress sensor and promotes cell survival through its dynamic interaction with hnRNPU and hnRNPA1. Oncogene 29, 1821–1834 (2010).

43. M. H. Wu, J. H. Chang, C. C. Chou, B. Y. Yung, Involvement of nucleophosmin/B23 in the response of HeLa cells to UV irradiation. Int. J. Cancer 97, 297–305 (2002).

44. T. Ukmar-Godec et al., Lysine/RNA-interactions drive and regulate biomolecular condensation. Nat. Commun. 10, 2909 (2019).

45. C. -D. Hu, Y. Chinenov, T. K. Kerppola, Visualization of interactions among bZIP and Rel family proteins in living cells using bimolecular fluorescence complementation. Mol. Cell 9, 789–798 (2002).

46. M. M. Bradford, A rapid and sensitive method for the quantitation of microgram quantities of protein utilizing the principle of protein-dye binding. Anal Biochem 72, 248–254 (1976).

47. T. L. Hwang, A. J. Shaka, Water suppression that works. excitation sculpting using arbitrary wave-forms and pulsed-field gradients. J. Magn. Reson. A. 112, 275–279 (1995).

48. W. -Y. Jeng, C. -Y. Chen, H. -C. Chang, W. -J. Chuang, Expression and characterization of recombinant human cytochrome *c* in E. coli. J. Bioenerg. Biomembr. 34, 423–431 (2002).

49. P. N. Palma, L. Krippahl, J. E. Wampler, J. J. Moura, BiGGER: A new (soft) docking algorithm for predicting protein interactions. Proteins: Struct., Funct., Bioinf. 39, 372–384 (2000).

50. E. F. Pettersen et al. UCSF Chimera—A visualization system for exploratory research and analysis. J. Comput. Chem. 25, 1605–1612 (2004).

51. W. Kabsch, XDS. Acta Crystallogr. D Biol. Crystallogr. 66, 125–132 (2010).

52. A. Vagin, A. Teplyakov, Molecular replacement with MOLREP. Acta Crystallogr. D Biol. Crystallogr. 66, 22–25 (2010).

53. P. D. Adams et al., PHENIX: a comprehensive Python-based system for macromolecular structure solution. Acta Crystallogr. D Biol. Crystallogr. 66, 213–221 (2010).

54. J. A. Mindell, N. Grigorieff, Accurate determination of local defocus and specimen tilt in electron microscopy. J. Struct. Biol. 142, 334–347 (2003).

55. J. M. De la Rosa-Trevín, et al., Scipion: A software framework toward integration, reproducibility and validation in 3D electron microscopy. J. Struct. Biol. 195, 93–99 (2016).

56. S. Chen, et al., High-resolution noise substitution to measure overfitting and validate resolution in 3D structure determination by single particle electron cryomicroscopy. Ultramicroscopy 135, 24–35 (2013).

57. P. A. Penczek, Three-dimensional spectral signal-to-noise ratio for a class of reconstruction algorithms. J. Struct. Biol. 138, 34–46 (2002).

58. Y. M. Liang et al., Novel nucleolar isolation method reveals rapid response of human nucleolar proteomes to serum stimulation. J. Proteomics 77, 521–530 (2012).

