## Supplementary Materials for "Mitochondrial cytochrome *c* liberates the nucleophosmin-sequestered ARF tumor suppressor in the nucleolus"

**This PDF file includes:**

Materials and Methods
Figures S1 to S13
Tables S1 and S2

### Supplementary Materials

### Materials and Methods

#### DNA constructs

For bimolecular fluorescence complementation (BiFC) experiments, the C*c*-encoding gene was fused with the C-end fragment of the yellow fluorescent protein (cYFP) in the cYFP vector, whereas the NPM-encoding gene was fused with the N-end fragment of the YFP in the nYFP vector. The cYFP and nYFP fragments were added to the C-terminal ends of C*c* and NPM, respectively. DNA for cloning was obtained via polymerase chain reaction (PCR) using the following primers: 5'-gtggaattcatgggtgatgttgagaaaggcaag-3' and 5'-cgagcggccgctcattagtagctttttgagat-3' for the C*c*-cYFP construct, and 5'-gtggaattcatggaagattcgatggacatggac-3' and 5'-cgaagcggccgcattttcttaagagacttcctcc-3' for the NPM-nYFP construct. The pBiFC-bJunYN155 and pBiFC-bFosYC155 plasmids were employed as positive controls, while the pBiFC-bJunYN155 and pBiFC-bFosΔZipYC155 plasmids were used as negative controls (*45*).

The gene encoding the native, full length NPM ($NPM_{(1\text{-}294)}$) was inserted into the pET28a(+) bacterial expression vector, along with an N-terminal His-tag. DNA for cloning was obtained by PCR using a cDNA sequence from Geneservice (UK). The oligonucleotides used in the PCR to obtain the $NPM_{(1\text{-}294)}$ construct were 5'- agccatatgatggaagattcgatggac-3' and 5'-gtgctcgagttaaagagacttcctcca-3'. The plasmid containing the truncated $NPM_{(9\text{-}122)}$ (*31*) species (cloned into the pET21a(+) vector) was acquired from Addgene. The plasmids for $NPM_{(225\text{-}294)}$, $NPM_{(123\text{-}294)}$ and $NPM_{(1\text{-}130)}$ were also cloned into the pET28a(+) vector. For PCR amplification, the following primers were used: 5'- agccatatggaatccttcaagaaacag -3' and 5'-gtgctcgagttaaagagacttcctcca-3' for $NPM_{(225\text{-}294)}$, 5'-cgcggcagccatatggcagagtcagaagatgaagag-3' and 5'- gtggtggtgctcgagttaaagagacttcctccactg- 3' for $NPM_{(123\text{-}294)}$, and 5'-agatgaagaggagtaggatgtgaaactc-3' and 5'-gagtttcacatcctactcctcttcatct-3' for $NPM_{(1\text{-}130)}$.

The genes coding for C*c* and yeast heme lyase for proper heme folding were cloned into the pBTR1 vector (*16*). The oligonucleotides employed to build the C*c* E104C mutant were 5'-gcgaccaattgctgatgaattc-3' and 3'-cgctggttaacgactacttaag-5'. All constructs were confirmed through automated sequencing.

#### Antibodies

Rabbit anti-human C*c* serum was obtained after immunizing male rabbits with recombinant C*c* suspended in a 0.85% solution of NaCl (20 mg/mL). The suspension was then incorporated into an equal amount of complete (Freund) adjuvant obtained from Difco (BD Biosciences). Secondary anti-rabbit IgG–FITC (fluorescein 5-isothiocyanate; catalog number F9887), mouse monoclonal anti-α-tubulin (α-tub, catalog number T8328), as well as secondary horseradish peroxidase (HRP)-conjugated anti-mouse IgG (catalog number A4416) and anti-rabbit IgG (catalog number A0545) were obtained from Sigma-Aldrich. Rabbit polyclonal antibody to COX-IV (cytochrome *c* oxidase, anti-COX IV; catalog number ab16056) and rabbit anti-histone H3 (catalog number ab1791) were from Abcam. Rabbit polyclonal anti-NPM was from Santa Cruz Biotechnology (catalog number sc-5564). Primary mouse anti-γ-H2AX (Ser139-phosphorylated histone H2AX) antibody (catalog number 05-636) was from EMD Millipore, while secondary mouse anti-rabbit IgG (conformation specific)-HRP conjugate (catalog number 5127S) was from Cell Signaling. Rabbit anti-eGFP (enhanced green fluorescent protein) was from BioVision (catalog number K817).

### Cell cultures and DNA damage induction

All cell lines were cultured at 37 °C in a humidified atmosphere of 5% $CO_2$. For immunofluorescence (IF) assays, HeLa and Cos-7 cells were cultured in Dulbecco's modified Eagle's medium (DMEM; Sigma-Aldrich) supplemented with 10% heat-inactivated fetal bovine serum (FBS; Sigma-Aldrich), 2 mM L-glutamine (Sigma-Aldrich), 100 U/mL streptomycin (Sigma-Aldrich) and 100 μg/mL penicillin (Sigma-Aldrich). MEF (mouse embryonic fibroblast) cells were cultured in DMEM medium with 4500 mg/L glucose (Sigma-Aldrich), supplemented with 10% FBS, 2 mM L-glutamine, 100 U/mL streptomycin, 100 μg/mL penicillin and 0.11 mg/mL sodium pyruvate (Sigma-Aldrich). For proteomic analysis, Jurkat human T-lymphoma cells were cultured in RPMI 1640 (PAA) supplemented as above. Jurkat cells were cultured as exponentially growing confluent monolayers in 645 mL flasks (Nunc) with the medium refreshed every 48 h. DNA damage was induced in Jurkat cells with 10 μM CPT (Sigma-Aldrich) for up to 6 h to ensure the release of C*c* from mitochondria into the cytosol. For BiFC assays, HEK293T cells were grown in DMEM supplemented as above. When used for fluorescence microscopy, the HEK293T cells were grown to 80% confluence in 24-well plates with 500 μL of DMEM, containing 20 mm glass coverslips. For subcellular fractioning and further immunoprecipitation (IP) analysis, HeLa cells were cultured in DMEM supplemented with the above components. HeLa cell aliquots (6 x $10^6$ cells) were seeded in 140 mm Petri dishes and treated with 20 μM CPT for the indicated incubation time. For nucleoli isolation, HeLa cells were grown to 80% confluence in in 140 mm Petri dishes with DMEM supplemented as above.

### Immunofluorescence

HeLa cells (30,000 per well) were grown for 24 h, whereas Cos-7 and MEF (7,500 per well) for 48 h, all of them on 20 mm glass coverslips, in 24-wells plates containing 500 μL of medium and treated with 20 μM CPT or 4 μM doxorubicin for 6 h. Then, 0.4 μM MitoTracker Red CMXRos (ThermoFisher Scientific) was added to medium and incubated for 20 min at 37 °C.

HeLa and MEF cells were fixed in a solution containing ethanol and acetic acid. For this, cells were washed once in phosphate buffer saline (PBS) and fixed for 20 min at –20 °C in a precooled solution of 95% ethanol and 5% acetic acid. Afterwards, cells were washed twice in PBS and treated with blocking buffer (10% FBS in PBS) for 30 min at room temperature (RT) and then incubated with rabbit anti-human C*c* serum (1/200 in blocking buffer) for 1 h. Coverslips were then washed three times with PBS for 5 min and probed with the secondary anti-rabbit IgG–FITC antibody (1/160 in blocking buffer) for 1 h, at RT, followed by three washes in PBS for 5 min.

Cos-7 cells were washed once in PBS and fixed for 40 min at RT in 4% paraformaldehyde (PAF, Sigma-Aldrich) prepared in 100 mM sodium phosphate buffer at pH 7.4. Then, cells were washed twice in PBS and permeabilized for 10 min at RT with 0.1% Triton X-100 prepared in 100 mM sodium phosphate buffer (pH 7.4). Afterwards, cells were washed once in PBS and incubated with rabbit anti-human C*c* serum (1/200 in 2% BSA; bovine serum albumin, Sigma-Aldrich, in PBS) for 1 h at RT. Coverslips were washed three times with PBS for 5 min and probed with the secondary anti-rabbit IgG–FITC antibody (1/160 in in 2% BSA-PBS) for 1 h at RT, followed by three washes in PBS for 5 min.

Alternatively, HeLa cells were also fixed in PAF. For this purpose, cells were washed once in PBS and fixed for 10 min at RT in 4% PAF prepared in PBS. Then, cells were washed twice in PBS and permeabilized for 5 min with precooled methanol at -20 °C. Afterwards, cells were washed once in PBS and blocked in a 3% BSA solution in PBS for 30 min at RT, and later incubated with rabbit anti-human C*c* serum (1/200 in 3% BSA-PBS) for 1 h. Coverslips were washed three times with PBS for 5 min and probed with the secondary anti-rabbit IgG–FITC antibody (1/160 in in 3% BSA-PBS) for 1 h at RT, followed by three washes in PBS for 5 min.

In all cases, nuclei were stained by incubation with Hoechst dye (Sigma-Aldrich; 200 μg/mL) for 10 min after the secondary antibody treatment, and cells were washed once with PBS. The slides were immersed in ethanol for 2 min, dried for a few minutes, mounted using n-propyl gallate (Sigma-Aldrich) and sealed with nail polish. Specimens were viewed using a Zeiss LSM 7 DUO scanning confocal microscope equipped with a plan-apochromat 63x/1.40 oil objective, a diode 405 nm, an argon 488 nm and a DPSS 561 nm lasers for viewing Hoechst, FITC and MitoTracker Red CMXRos, respectively.

**Cell extract preparation and protein purification by affinity chromatography**

Jurkat cell extracts from 1.8 L of culture with either untreated or 10 μM CPT-treated cells were prepared for affinity chromatography. Cells were harvested by centrifugation at 1,000 x g for 5 min, washed twice in PBS, pelleted again and resuspended to be lysed by sonication in buffer I (50 mM Tris-HCl [pH 7.5] with 50 mM NaCl and 0.25% Triton X-100) supplemented with 1 mM phenylmethylsulfonyl fluoride, 10 μg/mL aprotinin, 10 μg/mL leupeptin and 10 μg/mL of soybean trypsin inhibitor. Cellular debris was then removed by centrifugation at 20,000 x g for 30 min at 4 °C.

Affinity chromatography of resulting cell extracts was performed in a column with the E104C C*c* mutant covalently bound to thiol-Sepharose from Pharmacia (C*c* TS-4B), as previously described[1]. A blank column containing the Sepharose matrix alone, lacking C*c* (Blank TS-4B), was used as control. Jurkat T cell extracts, either untreated or treated with CPT, were thus loaded into the columns with and without C*c*. After loading, the columns were washed with 30 mL of buffer I (see above) and 30 mL of buffer II (50 mM Tris-HCl [pH 7.5] with 75 mM NaCl). Upon elution with 30 mL of buffer III (50 mM Tris-HCl [pH 7.5] with 300 mM NaCl), protein fractions were collected, lyophilized and stored at -80 °C.

**2D SDS-PAGE and protein preparation for mass spectrometry**

Protein samples resulting from affinity chromatography were analyzed using 2D SDS-PAGE. First, isoelectrofocusing (IEF) was carried out with the PROTEAN IEF Cell (Bio-Rad) system and 7 cm ReadyStrip IPG Strips (Bio-Rad) with linear pH gradients (pH 3-10), as described in Martínez-Fábregas et al. (2014). Afterwards, the IEF gels were loaded on a 12% polyacrylamide sodium dodecyl sulphate-polyacrylamide electrophoresis (SDS-PAGE) gel in a Mini-PROTEAN 3 Dodeca Cell (Bio-Rad). The 2D gels were stained using Blue Silver and analyzed using the PDQuest 2D Analysis Software Version 8.0.1 (Bio-Rad). Gel protein spots were manually excised using pipette tips. The selected proteins were reduced in-gel, alkylated and digested with trypsin as described (*14*). After digestion, the supernatant was collected and spotted onto a MALDI target plate and air-dried at room temperature. Later, 3 mg/mL of α-cyano-4-hydroxy-transcinnamic acid matrix (Sigma-Aldrich) in 50% acetonitrile was added to the dried peptide digested spots and then air-dried at room temperature.

**Matrix-Assisted Laser Desorption/Ionization Time-of-Flight Mass Spectrometry (MALDI-TOF MS)**

MALDI-TOF MS analyses were carried out in a 4800 Proteomics Analyzer MALDI-TOF/TOF mass spectrometer (Applied Biosystems) at the Genomics and Proteomics Center, Complutense University of Madrid. For protein identification, the UniProtKB-SwissProt database v.57.7 restricted to human protein (20,333 sequences) or NCBI database (10,084.244 sequences) were searched using a local license for MASCOT 2.1. Database search parameters and other details are mentioned in Martínez-Fábregas *et al.* (*14*).

**BiFC assays**

HEK293T cells were transiently transfected with the BiFC vectors and Lipofectamine 2000 Transfection Reagent (Invitrogen), following the manufacturer's instructions. 0.5 μg of DNA per construct was diluted into 50 μL of Opti-MEM medium (Invitrogen), 2 μL of Lipofectamine was diluted in 50 μL of Opti-MEM medium and the solutions were incubated separately. After a 5-min incubation at RT, the two solutions were mixed and incubated for other 20 min at RT. Finally, 100 μL of the DNA-Lipofectamine mixtures were added to the cells. To allow protein expression of both constructs, the cells were incubated at 37 °C for 24 h. Before imaging, DAPI (4', 6-diamidino-2-phenylindole) was added to cells at 500 ng/mL final concentration. Upon incubation at 37 °C for 30 min, untreated and CPT-treated HEK293T cells on coverslips were mounted in PBS, supplemented with 75% glycerol and observed under a Leica DM6000 B fluorescence microscope.

**Subcellular fractionation**

Separation of cell extracts into cytosol, membrane/organelle, nucleus and cytoskeleton fractions was performed with a ProteoExtract Subcellular Proteome Extraction Kit (Calbiochem), according to the manufacturer's indications. Purity of subcellular fractions was verified by Western blot analysis, using anti-α-tub, anti-COX IV and anti-histone H3 for specific detection of cytosol-, mitochondria- and nucleus-specific proteins, respectively.

**Western blot analysis**

For immunoblot detection of C*c*, α-tub, COX IV, histone H3 and NPM in the subcellular fractions or IP samples, protein content was determined using the DC™ protein assay (Bio-Rad Laboratories). For γ-H2AX immunodetection, total lysates were obtained after the addition of PBS buffer (supplemented with protease inhibitors) and sonication; protein content was determined with the Bradford Reagent (Bio-Rad). Proteins were resolved by SDS-PAGE in 12% gels and then transferred onto polyvinylidene fluoride (PVDF) membranes (EMD Millipore) using a Mini Trans-Blot electrophoretic transfer cell (Bio-Rad). Membranes were blocked in 5% non-fat dry milk in TBS (Tris-Buffered Saline) containing 1% Tween-20 (TPBS) and immunobloting was performed with primary antibodies. HRP-conjugated secondary antibodies were used for detection. The immunoreactive bands were detected using Amersham ECL Plus Western Blotting Detection Reagents (GE Healthcare Life Sciences) in a ChemiDoc™ Imaging System (Bio-Rad).

### Immunoprecipitation

Nuclear and cytosolic cell fractions from subcellular fractionation of control and CPT-treated HeLa cells were used for IP. Nuclear or cytosolic samples (300 μg protein) were incubated with 50 μL Sepharose 6B (Sigma-Aldrich) for 3 h at 4 ºC under stirring as a pre-clearing step to reduce nonspecific binding. Lysates were then centrifuged and 20 μL of rabbit anti-human C*c* was added to supernatants and incubated overnight at 4 ºC under stirring. As a negative control, 300 μg nuclear lysates were incubated with 1 μg anti-mouse immunoglobulin G (IgG) under the same conditions. Subsequently, 50 μL of Protein A Sepharose (GE Healthcare) was added to each lysate for 4 h at 4 ºC under stirring. The protein-Sepharose complexes were washed extensively, collected by centrifugation and boiled in freshly prepared reducing loading buffer. Controls including 30 μg of nuclear lysates were run concurrently with the IP samples in the Western blot assays.

### Protein expression and purification

Wild type (WT) C*c* or E104C C*c* mutant were produced in *E. coli* BL21(DE3) strains. 25 mL of overnight pre-cultures were shaken at 37 ºC in Luria-Bertani (LB) medium supplemented with 100 μg/mL ampicillin. 2.5 mL of pre-culture was used to inoculate 2.5 L of the same media in a 5 L Erlenmeyer flask. The bacterial culture was incubated at 30 ºC for 24 h, after which the cells were harvested at 6,000 rpm for 10 min using an Avanti J-25 refrigerated centrifuge (Beckman Coulter). Then, cells were resuspended in 1.5 mM borate buffer (pH 8.5), sonicated for 4 min and then centrifuged at 20,000 rpm for 20 min. For NMR measurements, $^{15}$N-labeled C*c* and $^{2}$H-$^{15}$N-labeled $NPM_{(1-130)}$ were produced in minimal media as previously described (*24*,*29*). $^{15}NH_4Cl$ was used as a nitrogen source.

In all cases, purification of recombinant C*c* from cell cultures was carried out as previously reported (*16*). For ITC measurements, the fractions containing C*c* were concentrated in an Amicon cell (3 kDa cut-off) until reaching the appropriate C*c* concentration and dialyzed against 10 mM sodium phosphate buffer (pH 7.4). For NMR titrations, C*c* was dialyzed against 5 mM sodium phosphate (pH 6.5).

NPM constructs were used to transform *E. coli* BL21(DE3) strains. Transformed cells with $NPM_{(1-294)}$, $NPM_{(225-294)}$, $NPM_{(123-294)}$ or $NPM_{(1-130)}$ vectors were harvested in fresh plates with 50 μg/mL kanamycin at 37 ºC, whereas cells expressing $NPM_{(9-122)}$ were grown in 100 μg/mL ampicillin. 250 mL pre-cultures in LB medium supplemented either with kanamycin or ampicillin were grown overnight and then used to inoculate 2.5-L cultures in 5-L flasks. After induction with 1 mM isopropyl β-D-1-thiogalactopyranoside (IPTG) and growth at 30 ºC for 24 h, the cells were harvested at 6,000 rpm for 10 min and resuspended in 40 mL lysis buffer composed of 20 mM Tris-HCl (pH 8.0), 0.8 M NaCl, 10 mM imidazole, 1 mM PMSF, 0.2 mg/mL lysozyme, 5 mM dithiothreitol (DTT) and 0.02 mg/mL DNase. The cell suspensions were sonicated for 4 min and centrifuged at 20,000 rpm for 20 min. Protein purification was achieved by affinity chromatography. Accordingly, the resulting cell lysates were loaded into a Ni Sepharose 6 Fast Flow column (GE Healthcare) previously equilibrated with lysis buffer. NPM proteins were eluted using an imidazole gradient from 0 to 300 mM. The protein-containing fractions were concentrated in an Amicon cell (10 kDa cut-off) until reaching the required protein concentration and dialyzed against 10 mM sodium phosphate buffer (pH 7.4) for ITC analysis or 5 mM sodium phosphate buffer (pH 6.5) for NMR assays. The purity of the protein samples was checked by SDS-PAGE analysis, and protein quantification was performed using the Bradford protein assay (*46*). With the only exception being $NPM_{(225\text{-}294)}$ and $NPM_{(123\text{-}294)}$, NPM molar concentration was expressed in relation to the pentameric form.

The synthetic peptide of mouse p19ARF herein used ($p19ARF_{(1\text{-}37)}$) comprised of 37 N-terminal residues was purchased from GeneCust. The peptide was weighted and reconstituted in the appropriate buffer without any observed aggregation.

**Circular dichroism**

CD (circular dichroism) spectra were recorded in the far-ultraviolet (UV) range (190-250 nm) at 25 ºC on a Jasco J-815 CD spectropolarimeter equipped with a Peltier temperature-control system, using a 1-mm quartz cuvette. In all cases NPM concentration was 3 μM, with the only exception being the $NPM_{(225\text{-}294)}$ construct (12 μM). Protein solutions were in 10 mM sodium phosphate buffer (pH 7.4). For each sample, 20 scans were averaged.

**EMSA**

$NPM_{(1\text{-}130)}$ (26 μM) was mixed with p19ARF (50 μM) and/or C*c* (12.5-100 μM) in 20 μL of 10 mM sodium phosphate buffer (pH 7.4), incubated for 30 min at RT and analyzed on a native 6% acrylamide gel in the same buffer at 100 V for 2 h. Proteins were stained with Coomassie Brilliant Blue.

**ITC titrations**

To analyze the interactions between C*c* and NPM constructs, ITC experiments were performed using a MicroCal Auto-iTC200 calorimeter (Malvern Instruments) at 25 ºC. The reference cell was filled with distilled water. The experiments consisted of successive 2-μL injections of a C*c* solution in 10 mM sodium phosphate buffer (pH 7.4) into the sample cell, which contained the different NPM constructs in the same buffer. The buffer for high-salt experiments contained 0.1 M KCl and 10 mM sodium phosphate, pH 7.4. Unless otherwise stated, ITC measurements were conducted with reduced C*c*. Protein concentrations were as follows: 300 μM C*c* and 20 μM $NPM_{(1\text{-}294)}$, 300 μM C*c* and 20 μM $NPM_{(225\text{-}294)}$, 200 μM C*c* and 10 μM $NPM_{(9\text{-}122)}$, and 300 μM C*c* and 10 μM $NPM_{(123\text{-}294)}$.

To analyze the interaction between the p19ARF peptide and the different NPM constructs, ITC measurements were carried out in a low volume Nano ITC (Waters-TA Instruments) at 25 ºC. The experiments consisted of successive 2-μL injections of p19ARF solution in 10 mM sodium phosphate buffer (pH 7.4) into the sample cell, which contained the different NPM constructs in the same buffer. Protein concentrations were as follows: 500 μM p19ARF peptide and 10 μM $NPM_{(1\text{-}294)}$, 250 μM p19ARF peptide and 10 μM $NPM_{(9\text{-}122)}$ complex. All solutions were degassed before titration. Titrant was injected at appropriate time intervals to ensure that the thermal power signal returned to the baseline prior to the following injection. To achieve homogeneous mixing in the cell, the stirring speed was maintained constant at 1,000 rpm. Experimental data – namely, the heat per injection normalized per mole of injectant vs. molar ratio – were analyzed with the Origin 7.0 software (OriginLab Corp.). Interaction models considering a single ligand binding site or two ligand binding sites, according to reference (*26*), were applied for estimating dissociation constants ($K_D$), enthalpies ($\Delta H$), and stoichiometries of interaction ($n$), from which the Gibbs energy ($\Delta G$) and the entropic contribution ($-T\Delta S$) for the interaction could be calculated. Because the general model for two ligand binding sites (*26*) indicated that neither cooperativity nor different binding sites could be observed for the interactions with stoichiometry 1:2, those were analyzed considering two identical and independent binding sites. Calibration and performance tests of the calorimeter were carried out by conducting $CaCl_2$–EDTA titrations with solutions provided by the manufacturer.

**SPR assays**

The formation of complexes between C*c* and $NPM_{(9-122)}$ or $NPM_{(1-294)}$ was assayed by SPR using a BiaCore 3000 and CM4 Chips. An automated desorption procedure was performed prior to each experiment to ensure the cleanliness of the BiaCore tubing, channels and sample injection port. The initial electrostatic attraction of C*c* to the CM4 Sensor Chip surface was assessed by considering its isoelectric point and was optimized to pH 5.8. The C*c* was then covalently attached to the matrix using standard amine-coupling chemistry. A reference flow cell was used as a control in which the chip surface was treated as described above, but without the injection of C*c*. The binding measurements were performed at 25 ºC using HBS-EP buffer containing 10 mM HEPES, 150 mM NaCl, 3 mM EDTA and 0.005 % surfactant P20, adjusted to pH 7.4. C*c* interactions were analyzed by flowing $NPM_{(9-122)}$ or $NPM_{(1-294)}$ at different concentrations (from 0.1 to 10 μM) over the C*c*-modified surface at a flow rate of 10 μL/min. Each concentration was injected at least three times. In each sensorgram, the signals from the reference flow cell surface were subtracted.

**NMR measurements**

1D $^1H$ NMR and 2D [$^1H$, $^{15}N$] HSQC spectra of C*c* was performed on a Bruker Avance 700 MHz at 25 ºC. Water signal was suppressed according to the excitation sculpting solvent suppression method (*47*). 1D $^1H$ spectra were recorded to monitor the Met80-methyl signal of reduced C*c* (13 μM) in the presence of 6 μM unlabeled $NPM_{(1-294)}$ or 26 μM $NPM_{(9-122)}$ and $p19ARF_{(1-37)}$ (40-100 μM) in 3 mm NMR tubes (0.25 ml volume). The synthetic peptide $p19ARF_{(1-37)}$ was weighted and reconstituted in 10 mM potassium phosphate buffer (pH 7) containing 1 mM TCEP without any observed aggregation.

C*c* interaction with $NPM_{(1-294)}$ and $NPM_{(9-122)}$ was followed by acquiring two-dimensional [$^1H$-$^{15}N$] HSQC spectra during titration of 50 μM $^{15}N$-labelled C*c* solutions with increasing amounts of each protein partner up to a final C*c*:NPM molar ratio of 1:0.25. Titration measurements were prepared in NMR tubes (Shigemi) up to a volume of 0.35 mL. 1D $^1H$ and 2D [$^1H$, $^{15}N$] HSQC measurements were made in 5 mM sodium phosphate buffer (pH 6.5).

Reverse titrations were performed by recording 2D [$^1H$-$^{15}N$] TROSY spectra of $^2H$-$^{15}N$-labeled $NPM_{(1-130)}$ at 250 μM, in the absence and in the presence of 500 μM non-labeled C*c* (C*c*:$NPM_{(1-130)}$ molar ratio of 2:1). In addition, TROSY spectra of 60 μM $^2H$-$^{15}N$-labeled $NPM_{(1-130)}$ mixed with 180 μM $p19ARF_{(1-37)}$ ($p19ARF_{(1-37)}$:$NPM_{(1-130)}$ molar ratio of 3:1) were also carried out. TROSY spectra were performed at the NMR Platform at Institut de Biologie Structurale (IBS, Grenoble, France) on a Bruker Avance 950 MHz at 25 ºC, and these measurements were made in 10 mM potassium phosphate buffer (pH 7) containing 1 mM TCEP.

All NMR measurements were made in the presence of 0.1 M sodium ascorbate to keep C*c* in its reduced state throughout the titration. pH values of the samples were verified after each titration step. To adjust the lock signal, 10% $D_2O$ was added. All NMR data were processed with the TopSpin NMR 2.0 software (Bruker), and the CSP (chemical-shift perturbation) analyses were performed with the Sparky 3 NMR Assignment Program (T.D. Goddard and D.G. Kneller, University of California – San Francisco, USA). The NMR signal assignments of the $^{15}$N and $^{1}$H nuclei of reduced C*c* (*48*) (BMRB accession number: 5406) and $NPM_{(1-130)}$ (*25*) (BMRB 19982) were already available.

**Molecular docking**

A soft docking algorithm implemented in the Biomolecular complex Generation with Global Evaluation and Ranking (BiGGER) software package (*49*) was used to generate *in silico* models of the C*c*:$NPM_{(9-122)}$ complex. For each run, 500 solutions were generated using a 15º angle step soft dock and a distance of 7 Å. Geometric docking solutions were generated based on the complementarity of protein surfaces. These solutions were evaluated and ranked according to their "global score", taking into account different interaction criteria including electrostatic energy of interaction, relative solvation energy and the relative propensity of side chains to interact. The center of mass for all structures was represented. CSP values were taken from NMR measurements and introduced as restraints in the docking calculations. The PDB files used for docking calculations were 1J3S for C*c* (*48*) and 5EHD for $NPM_{(9-122)}$. Results were evaluated with the zDOPE score, which ranged from -1.3 to -1.5. All molecular graphics of complexes were generated using the UCSF Chimera package (*50*).

**Crystallization, crystallographic data collection and structural determination**

The purified C*c:*$NPM_{(9-122)}$ complex was concentrated to 10 mg/mL in 20 mM HEPES buffer (pH 7.4). Good quality crystals were obtained in 25% PEG 400, 0.1 M MES buffer (pH 6.5) and 0.1 M $MgCl_2$, using the microbatch technique. Drops were prepared mixing 0.5 μL of protein with 0.5 μL of precipitant solution and 0.3 μL of additive 0.1 M TCEP hydrochloride. Crystals were cryoprotected in 30% v/v PEG 400 prior to vitrification.

Diffraction data were collected on the beamline bl13-Xaloc at the ALBA Synchrotron (Barcelona, Spain) with 0.25º oscillation range at 2.5 Å resolution. Data sets were processed using XDS (*51*) and scaled with AIMLESS from the CCP4 program suite. The C*c*:$NPM_{(9-122)}$ complex crystallized in the monoclinic $P2_1$ space group, with unit cell parameters $a$ = 58.8 Å, $b$ = 179.72 Å, $c$ = 104.3 Å, $\beta$ = 94.32 º. Asymmetric unit contains twenty monomers of the $NPM_{(9-122)}$ molecule.

The C*c*:$NPM_{(9-122)}$ complex structure was solved by the molecular replacement method using MOLREP (*52*) with the $NPM_{(9-122)}$ structure available at the PDB (PDB code 2P1B). Electron density maps were calculated using conventional 2Fo-Fc and Fo-Fc coefficients with the PHENIX suite (*53*). Intensive model building was performed with COOT. The refinement converged to the final values $R_{work}$ = 0.16 and $R_{free}$ = 0.23. No density was observed for the C*c* molecule in the complex. Data processing and refinement results are summarized in Table S2.

**Electron microscopy and image processing**

Aliquots of three samples ($NPM_{(1-294)}$, C*c*:$NPM_{(1-294)}$ and C*c*:$NPM_{(9-122)}$) were applied to 400 mesh grids (Maxtaform Cu/Rh HR26) coated with a thin (~8 nm) carbon layer and glow-discharged for 20 s. Grids were stained for 2 min with 2% uranyl acetate and air-dried before visualization. Images were acquired under minimal dose conditions with a FEI Tecnai G2 FEG200 electron microscope at 200 kV, at a nominal magnification of x50,000 and underfocus values ranging from -2 to -4 μm using a 16-megapixel FEI Eagle CCD (the pixel size of the acquired images was 2.16 Å). The contrast transfer function of each image was estimated using the CTFFIND3 program (*54*). Micrographs with visible drift and astigmatism were discarded. Single particles were selected manually, extracted from micrographs and normalized using Scipion (*55*), a suite that integrates several software packages. Particles were classified using a free-pattern maximum-likelihood method (Relion 2D-Classification). To evaluate the structural homogeneity of the different data sets, Relion 3D classifications were performed (*56*). The final reconstruction step of the three samples was performed by projection matching, using C5 symmetry. Low-resolution of the final 3D models of $NPM_{(1-294)}$, C*c*:$NPM_{(1-294)}$ and C*c*:$NPM_{(9-122)}$ complexes was estimated to be 22.5 Å, 23.5 Å and 25 Å, respectively, based on the FSC criterion (*57*) (Fourier shell correlation). Chimera (*50*) was used for visualization of the volumes and docking of the atomic structures.

**Fluorescent labeling**

$NPM_{(1-294)}$ (20 μM) was incubated with 200 μM maleimide Oregon Green 488 (Molecular Probes) to label its exposed Cys104. $p19ARF_{(1-37)}$ (100 μM) was labeled in its Lys26 with 350 μM NHS-Rhodamine (Thermo Scientific), whereas C*c* E104C mutant (50 μM) in Cys104 with 125 μM Alexa Fluor 647 (Molecular Probes). Protein labeling was performed for 2 h at RT in the dark, in 10 mM sodium phosphate buffer at pH 7.4, containing 100 mM NaCl. Labeling procedures were achieved according to the manufacture's protocol. Then, labeled proteins were purified in PD-10 columns (GE Healthcare) and concentrated in Amicon® Ultra 4 mL centrifugal filters (30 kDa or 3.5 kDa cut-off for NPM or p19ARF/C*c*, respectively; Merck), until reaching the required protein concentration. Protein concentration was measured by recording UV-Vis spectra and following the manufacture's indications.

**Phase separation assays**

Non-labelled $NPM_{(1-294)}$ (5 μM) was prepared in either 10 mM Tris-HCl buffer, 100 mM NaCl, 2 mM DTT, pH 7.5 or in 10 mM sodium phosphate, 100 mM KCl, pH 7.4, in a total solution volume of 20 μL. For droplet formation, non-labelled $p19ARF_{(1-37)}$ or non-labelled C*c* were added at 5:1 and 10:1 ratios for the complexes $p19ARF_{(1-37)}$:$NPM_{(1-294)}$ and C*c*:$NPM_{(1-294)}$. The samples (2 μL) were imaged in sealed chambers comprised of glass slide and coverslip kept together by a layer of 3M 300 LSE high-temperature double-sided tape. The samples were equilibrated at RT for 15 min before imaging. DIC images were taken using an Automated Inverted Microscope TIRF - ScanR Olympus with a Hamamatsu Orca-ER camera and a 60x/1.20 water UPlan SAPo objective.

For fluorescence droplet analysis, we mixed 5% fluorescently labeled with 95% unlabeled proteins. Thus, 5 μM $NPM_{(1-294)}$ was mixed with either 75 μM $p19ARF_{(1-37)}$ or 75 μM E104C C*c* mutant in 10 mM Tris-HCl buffer, 100 mM NaCl, 2 mM DTT, pH 7.5. For ternary mixture assays, 5 μM $NPM_{(1-294)}$ was mixed with 50 μM $p19ARF_{(1-37)}$ and then titrated with increasing concentrations of E104C C*c* mutant. Fluorescent images were taken in a Zeiss LSM780 confocal microscope system, equipped with an Ar-laser (458, 488, 514 nm); HeNe-laser (543 nm) 1 mW; DPSS-laser (561 nm) 20 mW; HeNe-laser (594 nm) 2 mW; HeNe-laser (633 nm) 5 mW. The samples (30 μL) were placed in multiwall plates and imaged using a 63x oil objective. The samples were equilibrated at RT for 15 min before imaging.

**Nucleoli isolation**

Method for isolating nucleoli was conducted as published (*58*). One hour before nucleolar isolation, medium of HeLa cells was replaced with fresh, pre-warmed DMEM. Then, the culture medium was decanted from the dishes, and cells were washed with 10 mL cold solution I (0.5 M sucrose, 3 mM $MgCl_2$ and Roche's Complete Protease Inhibitor Cocktail). The solution was quickly decanted, followed by 2 quick rinses with the same volume of solution I. Cells were scraped from the plate on ice and placed into a 15 mL tube. Solution I was added into the collected cells to make up a volume of 3 mL. To break down the cells and release the nucleoli, the cells were sonicated once on ice at 30% amplitude, 5 s on, 30 s off. The isolated nucleoli were further incubated on ice in 1.5 mL Eppendorf tubes with 5 μM Oregon Green 488-$NPM_{(1-294)}$, 5 μM Rhodamine-$p19ARF_{(1-37)}$ and Alexa Fluor 647-C*c* (from 2.5 to 20 μM) for 30 min or the specified period of time. Prior to observation, tubes were centrifuged at 1800 ×g for 5 min at 4 °C, and the resulting pellet containing isolated nucleoli was resuspended in cold solution I. As nucleoli are active fluids requiring metabolism of ATP to maintain this fluidity, ATP was not depleted from the nucleolar preparations. Nucleoli were imaged in a Leica TCS SP2 laser-scanning microscope (Leica Microsystems) with a 63x oil immersion objective. Samples were excited with an Ar-laser at 488 nm, a HeNe-laser at 543 nm and a HeNe laser at 633 nm. Fluorescence emission was detected at 510-540 nm (green), 560-600 nm (red) or 670-750 nm (far red).

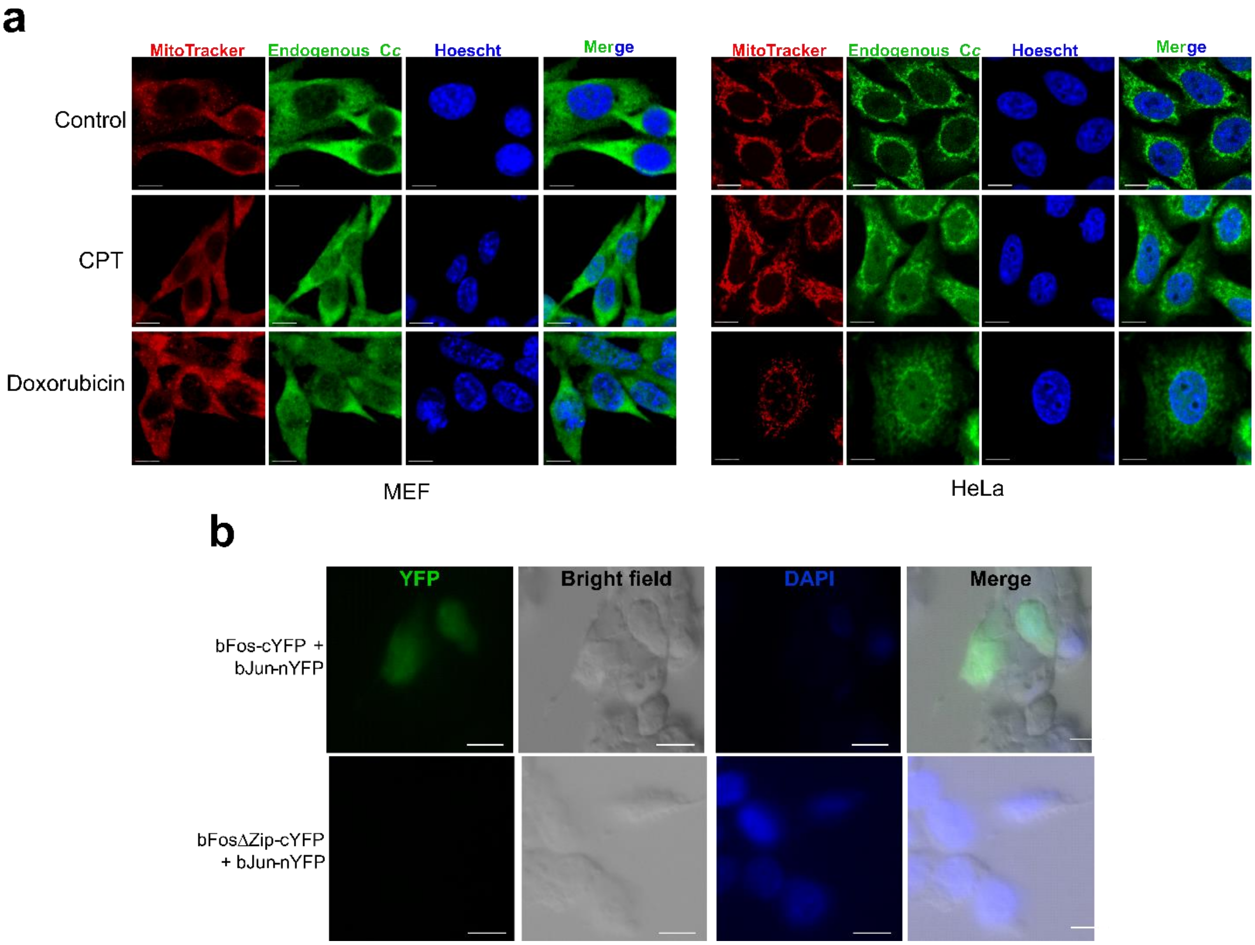


**Fig. S1. | a,** Immunofluorescence analysis of endogenous C*c* in MEF (*left*) and HeLa (*right*) cells, upon treatment with 20 μM CPT or 4 μM doxorubicin for 6 h. C*c* was visualized with an anti-C*c* antibody (green fluorescence) using a confocal microscope (x63 oil objective). Mitochondria were stained with MitoTracker Red CMXRos (red fluorescence) and nuclei with Hoechst (blue). Non-treated cells were used as control. Co-localization of green C*c* fluorescence and blue nuclear staining is shown in the merge images. Scale bars are 10 μm. HeLa cells were fixed in 4% PAF, whereas MEF were fixed in a 95% ethanol and 5% acetic acid solution. **b,** Positive and negative controls for BiFC assays. HEK293T cells were transfected with the N-terminal YFP fragment (nYFP) attached to bJun (bJun-nYFP), along with the bFos-cYFP vector containing the C-end domain of YFP bound to bFos (positive control, *upper*). For negative controls, a construct containing bFos lacking the bJun-interacting domain (bFosΔZip) was used (*lower*). Reconstruction of the YFP protein leads to fluorescence emission. Scale bars are 5 μm.

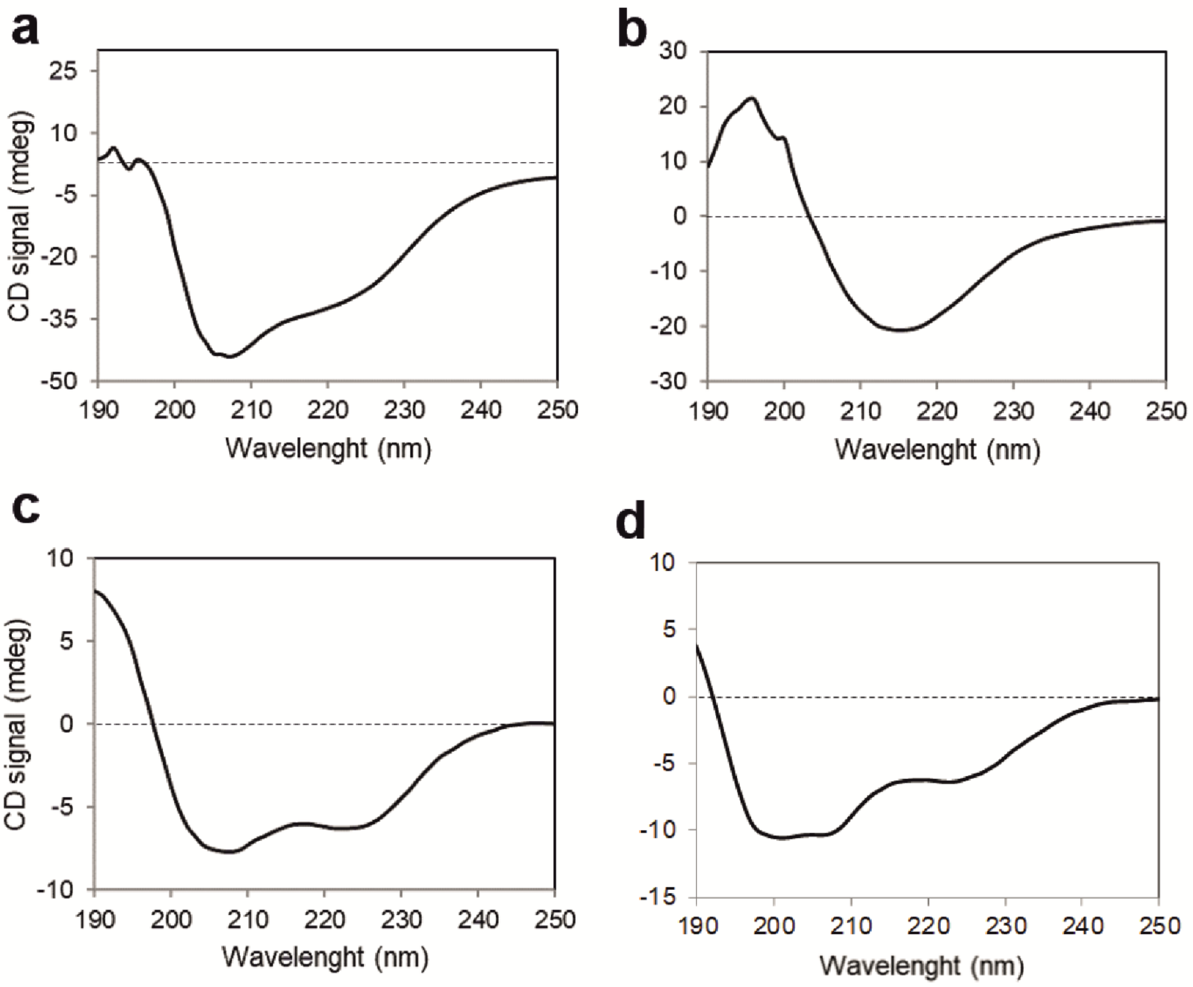


**Fig. S2 |** Far-UV CD spectra of NPM constructs NPM$_{(1-294)}$ (**a**), NPM$_{(9-122)}$ (**b**), NPM$_{(225-294)}$ (**c**) and NPM$_{(123-294)}$ (**d**).

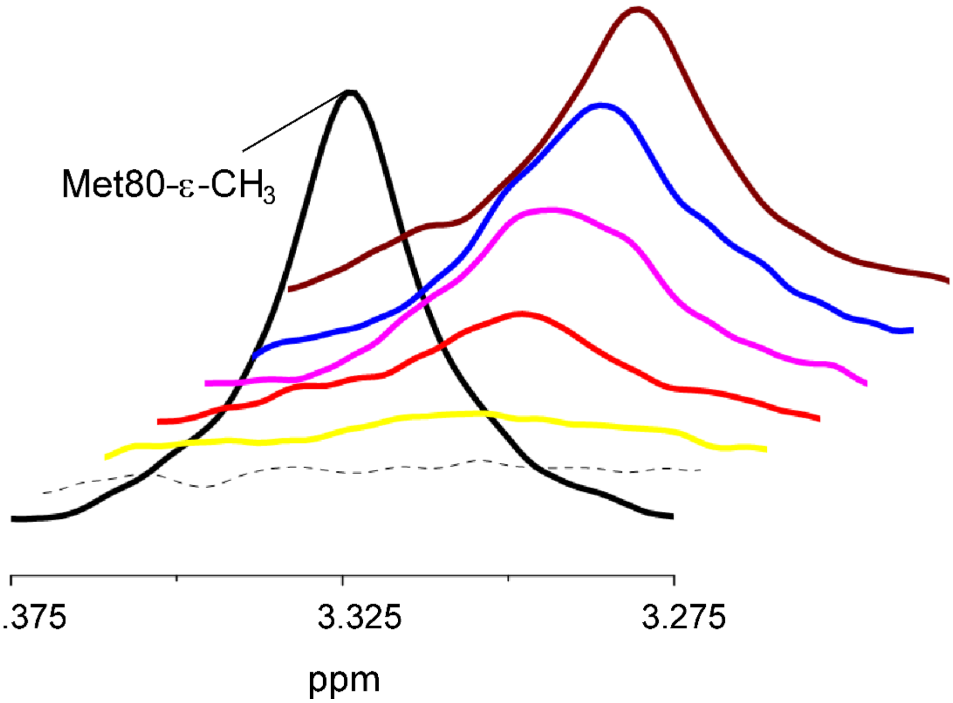


**Fig. S3 | Competition between C*c* and p19ARF for NPM(9-122) binding.** 1D [1]H NMR spectra monitoring the Met80-methyl signal of 13 μM reduced C*c*, either free (solid black) or upon successive additions of 26 μM NPM(9-122) (dotted line) and the p19ARF peptide at increasing concentrations of 40 μM (yellow), 50 μM (red), 52 μM (magenta), 55 μM (blue) and 57 μM (brown).

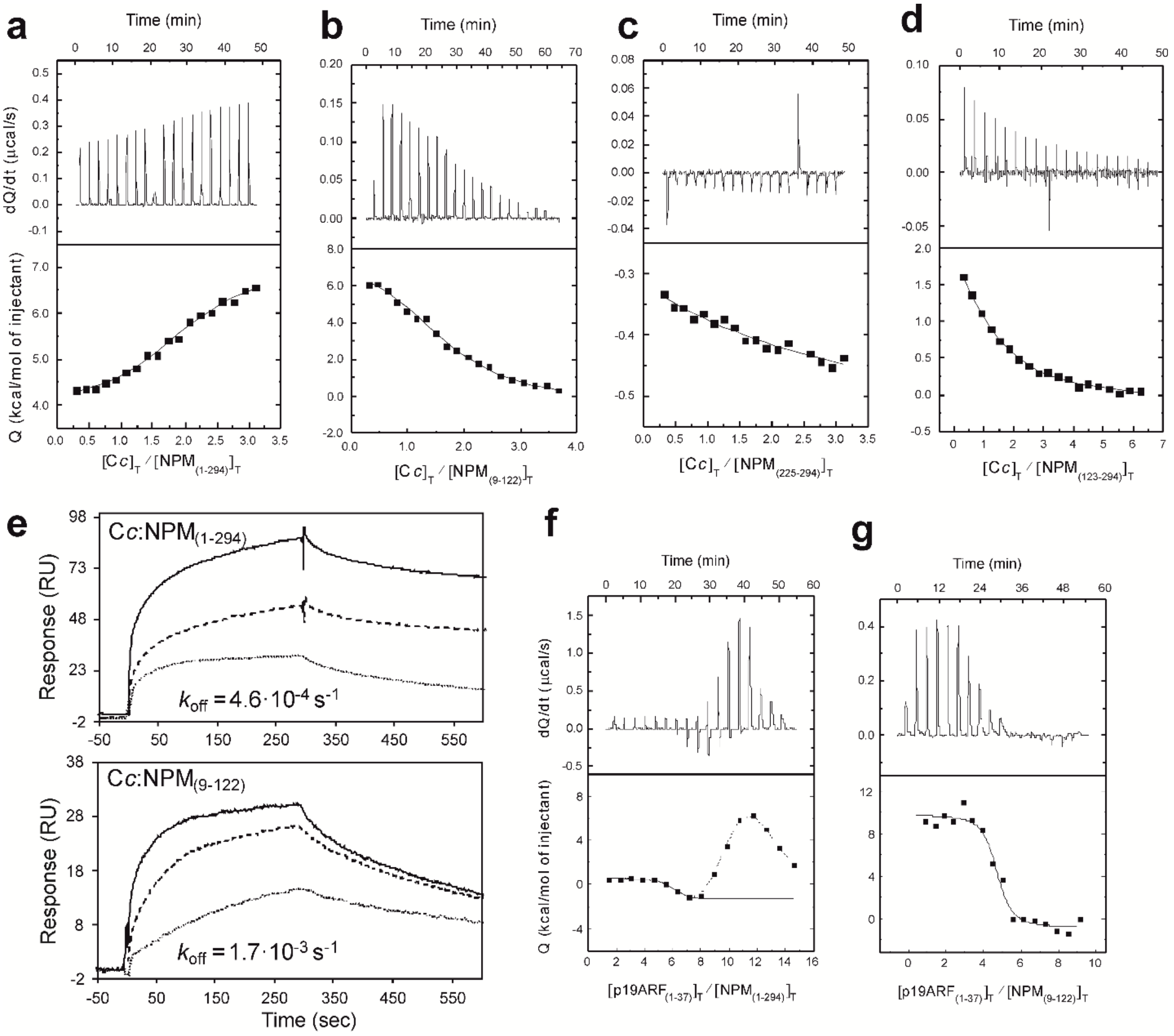


**Fig. S4 | Thermodynamic and kinetic analysis of the C*c* and p19ARF interaction with NPM constructs. a-d,** ITC titrations of C*c* with $NPM_{(1-294)}$ (**a**), $NPM_{(9-122)}$ (**b**), $NPM_{(225-294)}$ (**c**) and $NPM_{(123-294)}$ (**d**). Thermograms (*upper plots*) and binding isotherms (*lower plots*) are shown. **e,** SPR sensorgrams for the binding of C*c* to $NPM_{(1-294)}$ (*upper plot*) or $NPM_{(9-122)}$ (*lower plot*). $NPM_{(1-294)}$ concentrations were 0.1, 0.5 and 1.0 μM, whereas $NPM_{(9-122)}$ concentrations were 1, 5 and 10 μM. Three replicate injections were performed for each protein concentration. In each sensorgram, the signals from the control surface were subtracted. Kinetic dissociation constant ($k_{off}$) values are shown. **f-g,** ITC titration of the p19ARF peptide with $NPM_{(1-294)}$ (**f**) and $NPM_{(9-122)}$ (**g**). Thermograms (*upper plots*) and binding isotherms (*lower plots*) are shown.

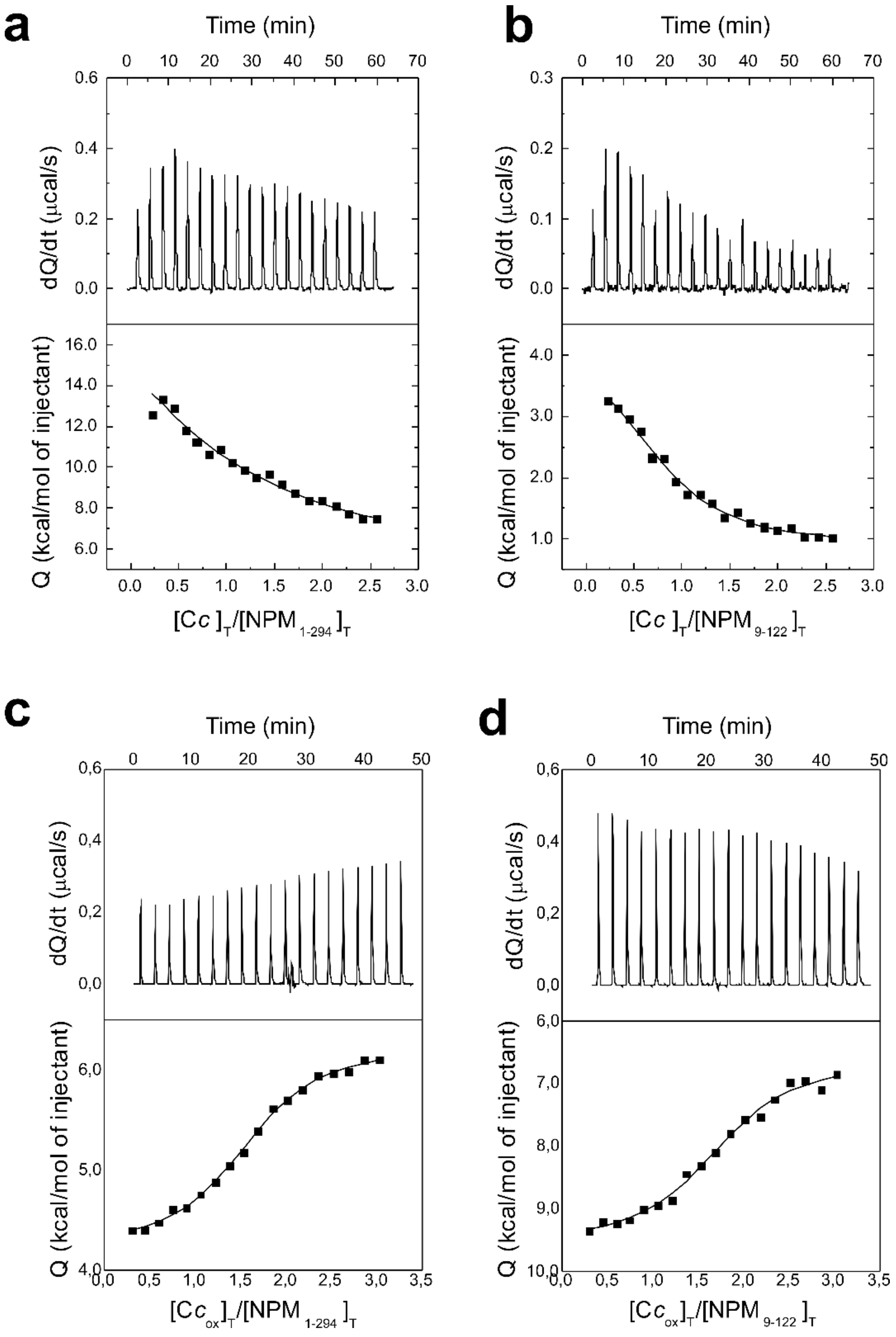


**Fig. S5 | a-b, ITC titrations of C*c* with NPM$_{(1-294)}$ and NPM$_{(9-122)}$ in the presence of 0.1 M KCl. c-d, ITC titrations of oxidized C*c* with NPM$_{(1-294)}$ and NPM$_{(9-122)}$.** Thermograms (*upper*) and binding isotherms (*lower*) are shown.

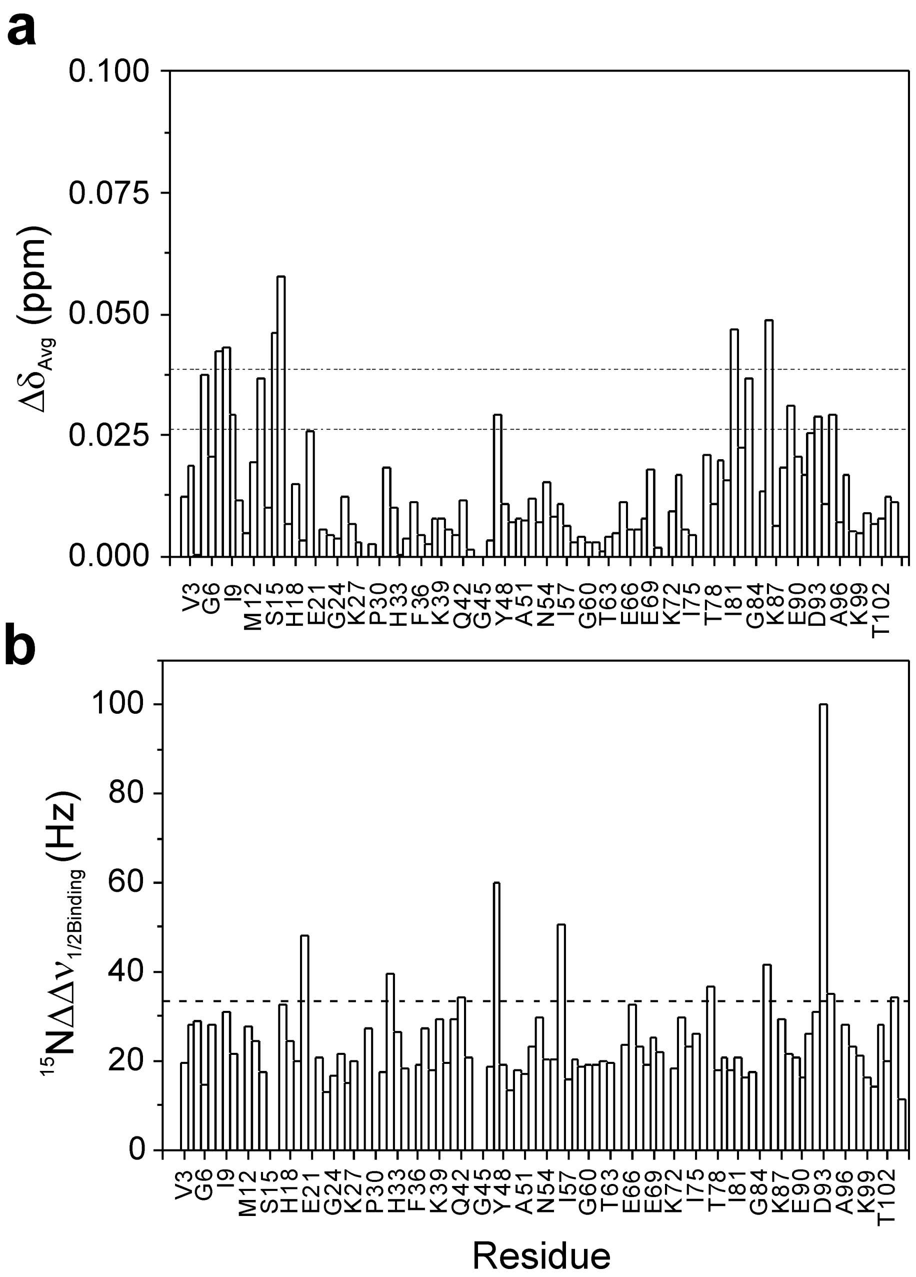


**Fig. S6 | NMR titrations of $^{15}$N-labeled C*c* with NPM$_{(9\text{-}122)}$.** ***Upper,*** Average CSPs ($\Delta\delta_{Avg}$) experienced by amide resonances of C*c* in complex with NPM$_{(9\text{-}122)}$ at the C*c*:NPM ratio of 1:0.5. ***Lower,*** Differences in the line-width ($\Delta\Delta\nu_{1/2Binding}$) of $^{15}$N dimension of the NMR amide signals of C*c* upon binding to NPM$_{(9\text{-}122)}$. The $\Delta\Delta\nu_{1/2Binding}$ values were calculated from the difference between [$^{1}$H-$^{15}$N] HSQC spectra of free C*c* and those of the complex at the above molar ratio. The threshold (dashed line) corresponds to the average plus two times the standard deviation ($\Delta\Delta\nu_{1/2Binding} \geq \Delta\Delta\nu_{1/2Binding}$ + 2Sn-1). Residues beyond the threshold exhibit significant line broadening.

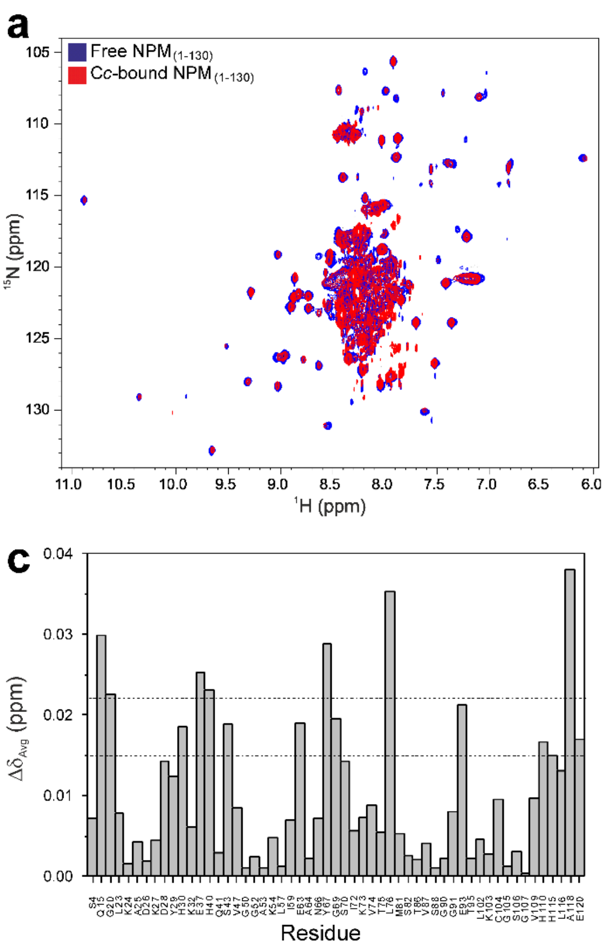

**Fig. S7 | NMR titrations of $^{2}H$-$^{15}N$-labeled NPM$_{(1\text{-}130)}$ with C*c* at the C*c*:NPM$_{(1\text{-}130)}$ molar ratio of 2:1 (a) or p19ARF$_{(1\text{-}37)}$ at the p19ARF$_{(1\text{-}37)}$:NPM$_{(1\text{-}130)}$ molar ratio of 3:1 (b)**. Free and bound NPM$_{(1\text{-}130)}$ spectra are represented in blue and red, respectively. **c-d**, Average CSPs ($\Delta\delta_{Avg}$) experienced by amide NMR signals of NPM$_{(1\text{-}130)}$ after binding to C*c* (**c**) and p19ARF$_{(1\text{-}37)}$ (**d**) determined at the indicated molar ratios. The threshold (dashed line) corresponds to the average plus two times the standard deviation ($\Delta\Delta\nu_{1/2Binding} \geq \Delta\Delta\nu_{1/2Binding}$ + 2Sn-1).

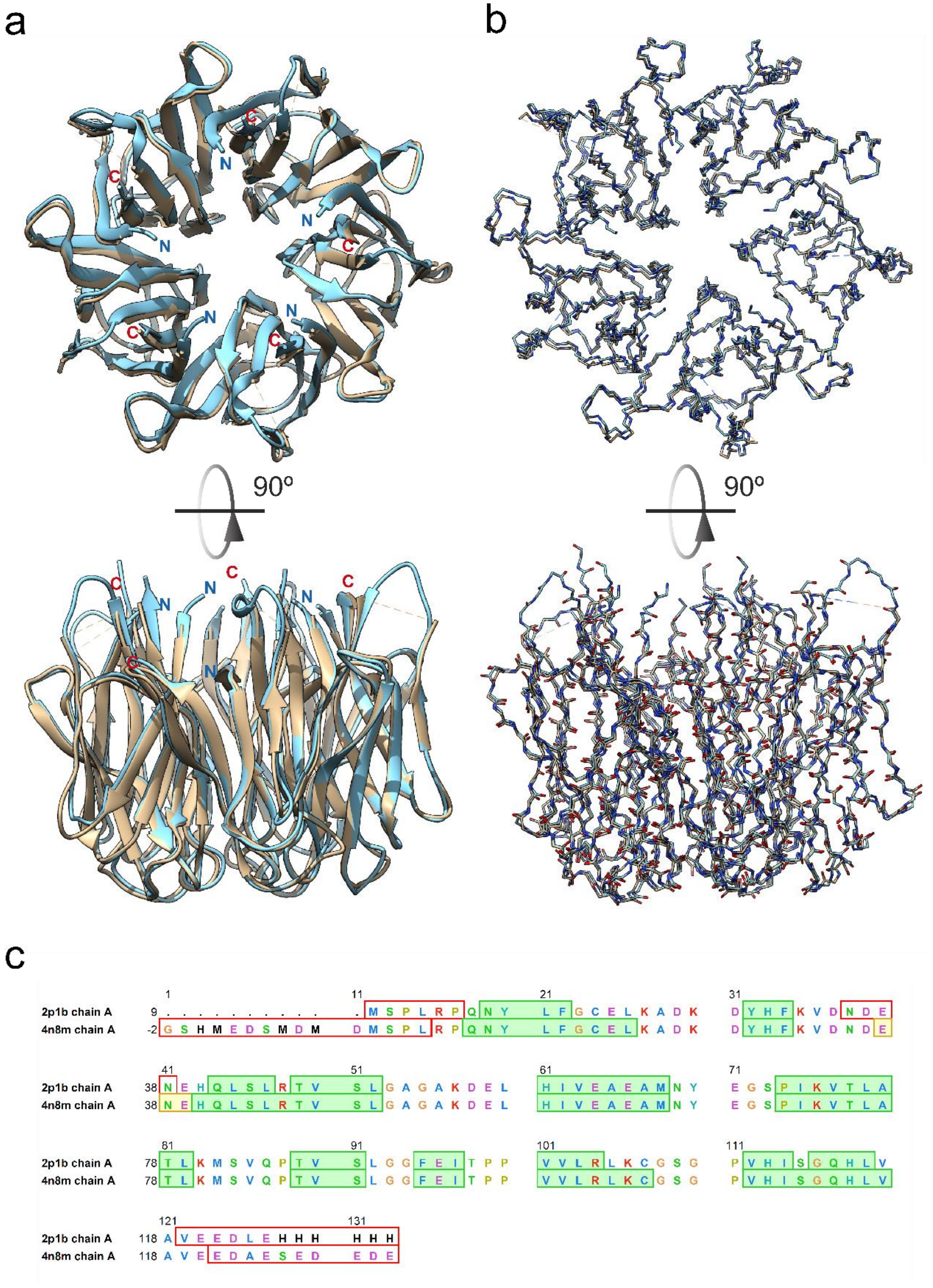


**Fig. S8 | Overlay of the structures of NPM core with (PDB code 2P1B; beige; *35*) and without (PDB code 4N8M; cyan; *25*) a 6x-His tag at the C-end. a,** Richardson's ribbon diagrams. **b,** Full backbone and main-chain atoms in the upper and lower views, respectively. **c,** A snapshot of the sequence alignment of the two proteins as shown in UCSF Chimera. Green shadows highlight β strands. Pale yellow background corresponds to helical (α) regions. Red empty boxes remark regions for which structural data is missing in the PDB files. Residue symbols are colored according to the Clustal-X pattern.

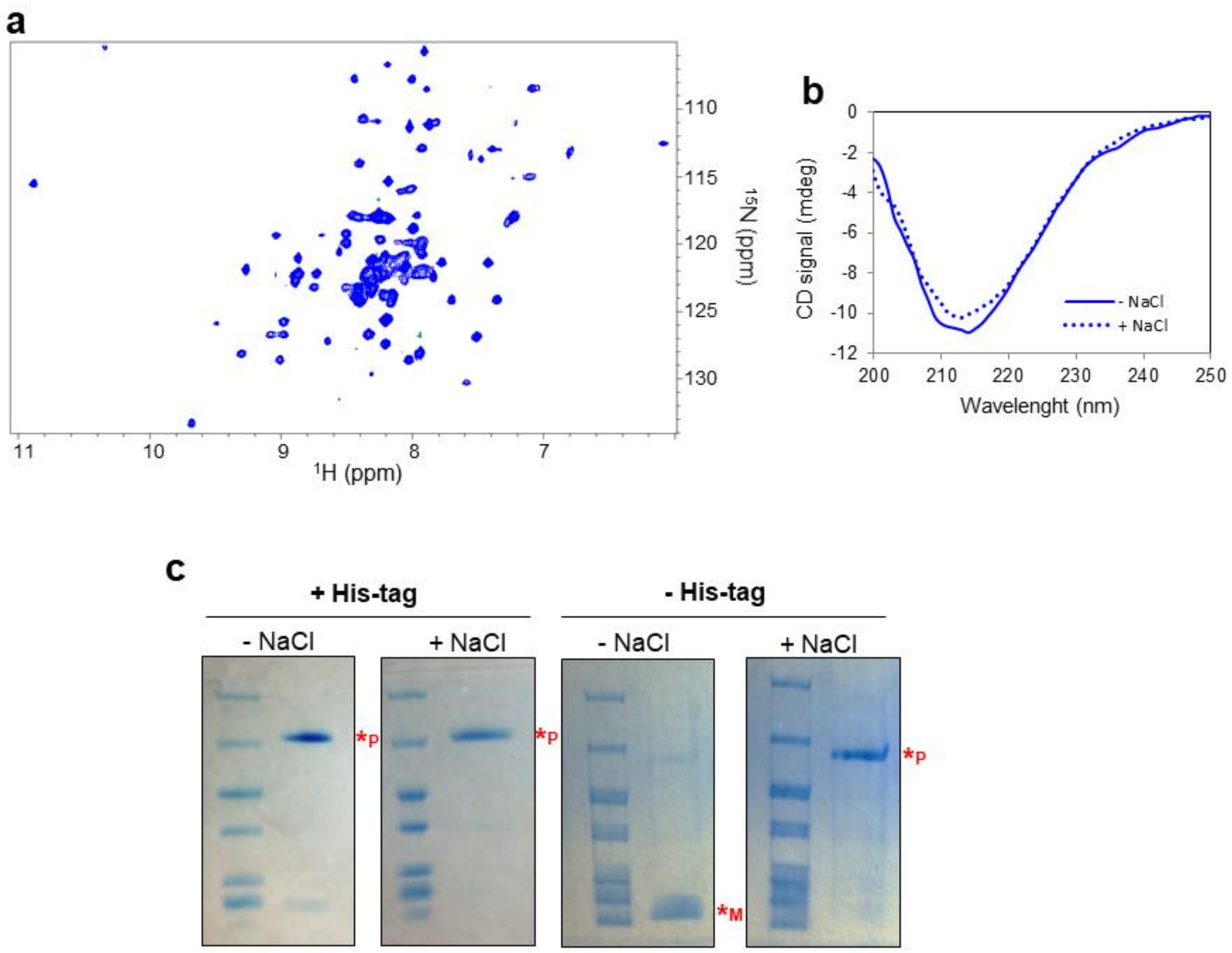


**Fig. S9 | a,** Two-dimensional transverse relaxation-optimized spectroscopy (2D TROSY) spectrum of His-tagged $^{2}$H-$^{15}$N-labeled NPM$_{(1\text{-}130)}$ (50 µM) in 10 mM potassium phosphate pH 7.0 containing 1 mM TCEP, as recorded at a Bruker Avance 700 MHz NMR spectrometer at 298 K. **b,** CD spectra of 3 µM His-tagged NPM$_{(1\text{-}130)}$ in 10 mM potassium phosphate pH 7.0 with or without 150 mM NaCl. **c,** SDS-PAGE of non-labelled NPM$_{(1\text{-}130)}$ with (*left*) or without (*right*) a His-tag, under low salt (-NaCl) or high salt (+NaCl) concentrations. Those bands matching the molecular mass of the pentamer or monomer are indicated as *P or *M, respectively. Molecular weight markers are on the left lane at each panel.

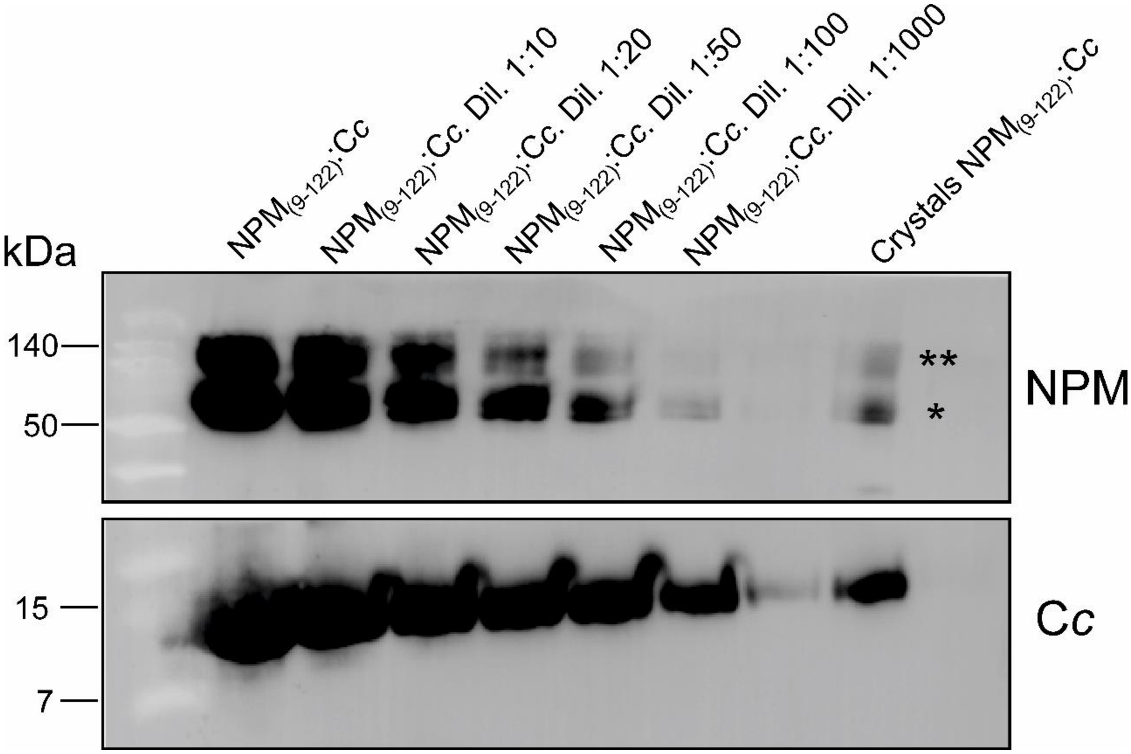


**Fig. S10 |** Immunoblotting against NPM and C*c* in the non-diluted complex $NPM_{(9\text{-}122)}$:C*c*, the diluted $NPM_{(9\text{-}122)}$:C*c* complexes (ranging from 1:10 to 1:1000) and in the extensively washed crystals of the same complex. One asterisk stands for the NPM pentamer (67.6 kDa), whereas two asterisks indicate an NPM decamer (135.2 kDa).

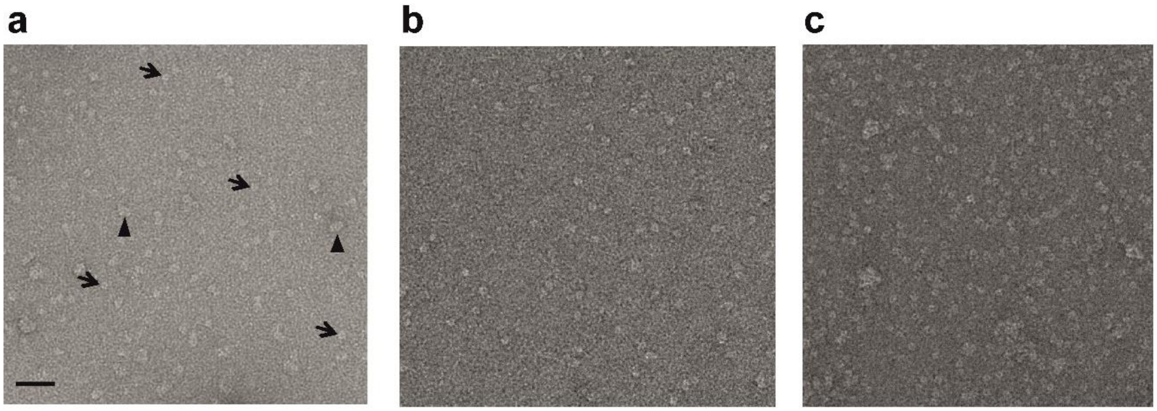

**Fig. S11 | Negatively stained electron micrographs of NPM$_{(1\text{-}294)}$ (a), C*c*:NPM$_{(1\text{-}294)}$ (b) and C*c:*NPM$_{(9\text{-}122)}$ (c),** from which three-dimensional reconstructions were carried out. In **a**, front and side views are indicated by arrows and arrowheads, respectively. Scale bar is 50 nm.

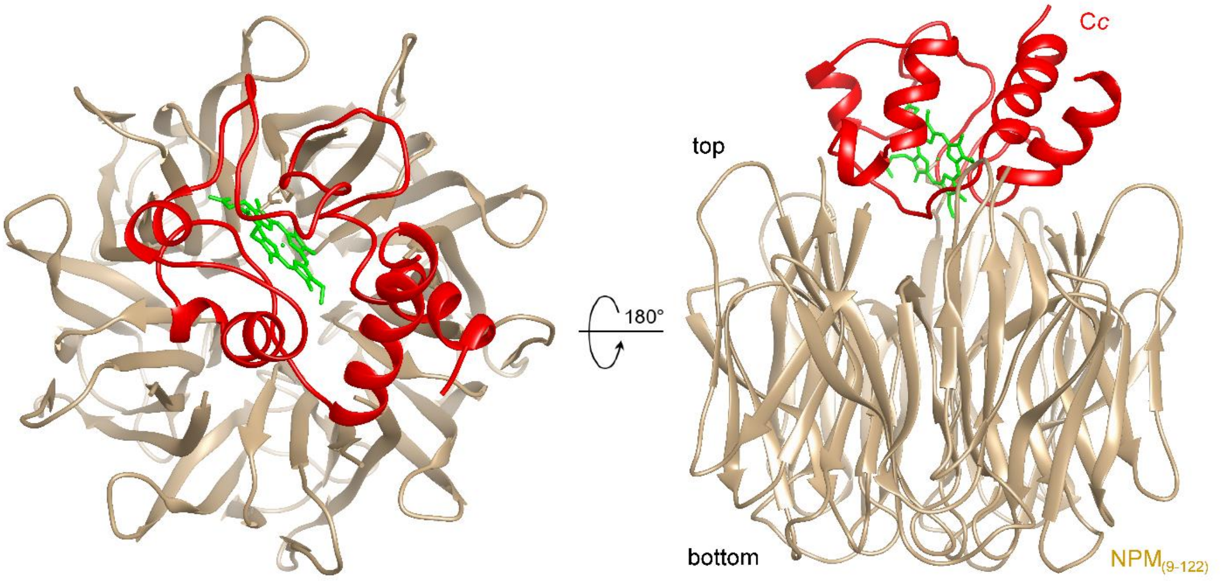


**Fig. S12 | NMR-based BiGGER molecular docking of the C*c*:NPM$_{(9\text{-}122)}$ complex.** Ribbon representation corresponds to the best complex model inferred from global calculation scores. NPM$_{(9\text{-}122)}$ and C*c* ribbons are shown in beige and red respectively, the C*c* heme group is in green. The structural PDB coordinates of NPM$_{(9\text{-}122)}$ (PDB code 5EHD) and C*c* (PDB code 1J3S) were used as inputs for docking calculations.

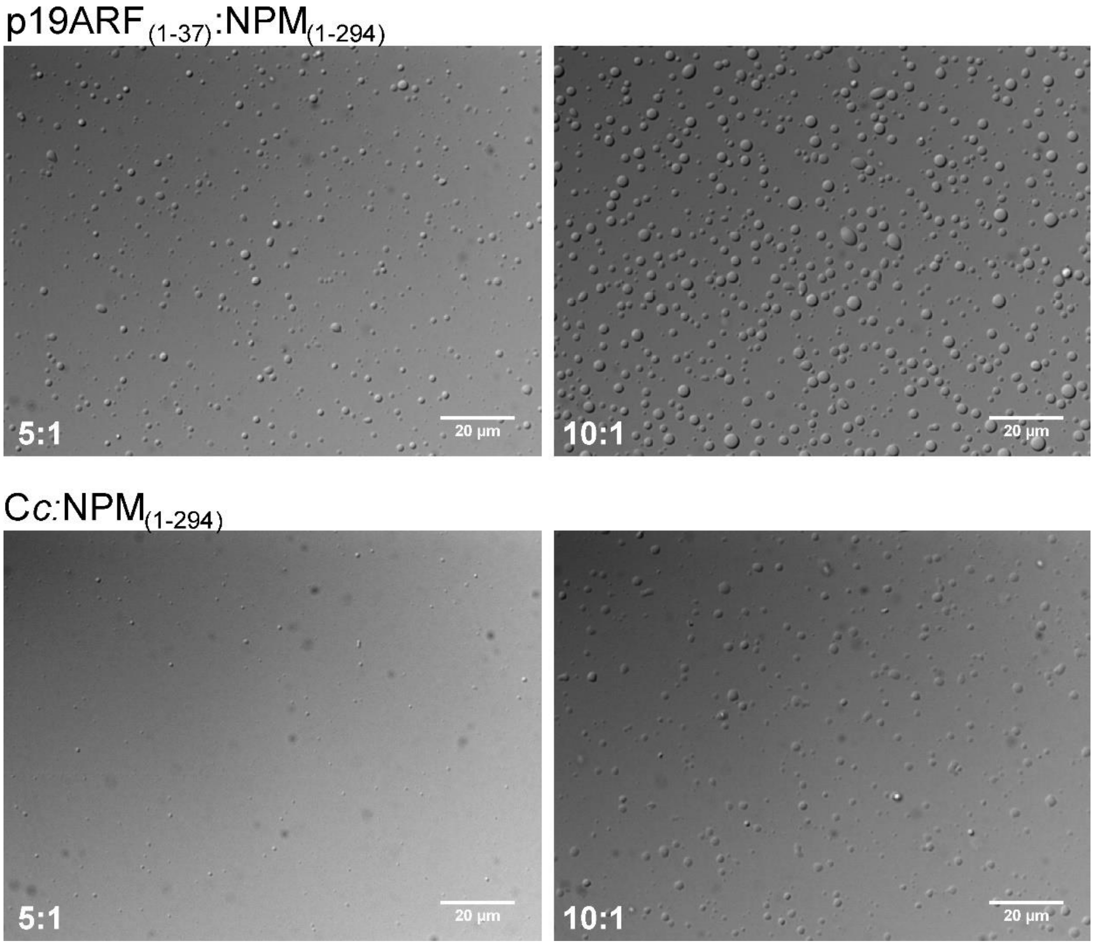


**Fig. S13 | *In vitro* droplets formed by NPM$_{(1\text{-}294)}$ with either p19ARF or C*c*** in 10 mM sodium phosphate buffer (pH 7.4) containing 100 mM KCl. The DIC microscopy images show phase separation induced by NPM$_{(1\text{-}294)}$ with p19ARF$_{(1\text{-}37)}$ (*upper panel*) or C*c* (*lower panel*). NPM$_{(1\text{-}294)}$ was used at 5 μM, whereas p19ARF$_{(1\text{-}37)}$ and C*c* were added at p19ARF$_{(1\text{-}37)}$:NPM$_{(1\text{-}294)}$ and C*c*:NPM$_{(1\text{-}294)}$ ratios of 5:1 and 10:1.

**Table S1. Thermodynamic values inferred from ITC measurements**

Equilibrium thermodynamic parameters for the interactions between the NPM constructs and C*c* and p19ARF$_{(1\text{-}37)}$ in 10 mM sodium phosphate buffer at pH 7.4. Gibbs free energy ($\Delta G$), enthalpy ($\Delta H$), entropic term ($-T\Delta S$), equilibrium dissociation constant ($K_D$), and stoichiometry of the reaction ($n$) are shown. The affinity of a protein-protein interaction is defined by the Gibbs energy of the binding: $\Delta G = -RT \ln K_A = RT \ln K_D$. $\Delta G$ has two different contributions, $\Delta H$ and $-T\Delta S$, according to the equation: $\Delta G = \Delta H - T\Delta S$. NPM concentration refers to the pentameric form in all cases, but to the monomeric form in the case of the NPM$_{(123\text{-}294)}$ construct. *Parameters measured in 10 mM sodium phosphate buffer at pH 7.4 containing 0.1 M KCl. **Parameters measured with oxidized C*c*. ***Stoichiometry value fixed to 1 in order to guarantee convergence because of the low binding affinity.

| **Protein complex** | **$\Delta G$ (kcal/mol)** | **$\Delta H$ (kcal/mol)** | **($-T\Delta S$) (kcal/mol)** | **$K_D$ (μM)** | ***n*** |
|---|---|---|---|---|---|
| C*c*:NPM$_{(1\text{-}294)}$ | -7.3 | -3.1 | -4.2 | 4.20 | 2.0 |
| C*c*:NPM$_{(9\text{-}122)}$ | -7.1 | 8.1 | -15.2 | 5.90 | 1.7 |
| C*c*:NPM$_{(225\text{-}294)}$ | -5.0 | 2.7 | -7.7 | 204.1 | 1*** |
| C*c*:NPM$_{(123\text{-}294)}$ | -6.7 | 4.3 | -11.0 | 13.0 | 1.0 |
| p19ARF$_{(1\text{-}37)}$:NPM$_{(1\text{-}294)}$ | -8.9 | 1.9 | -10.8 | 0.28 | 5.6 |
| p19ARF$_{(1\text{-}37)}$:NPM$_{(9\text{-}122)}$ | -8.9 | 14.7 | -23.6 | 0.29 | 4.4 |
| C*c*:NPM$_{(1\text{-}294)}$* | -5.8 | 19.8 | -25.6 | 53.0 | 1.1 |
| C*c*:NPM$_{(9\text{-}122)}$* | -6.5 | 3.4 | -9.9 | 18.0 | 0.8 |
| C*c*:NPM$_{(1\text{-}294)}$** | -7.7 | -2.0 | -5.7 | 2.27 | 1.6 |
| C*c*:NPM$_{(9\text{-}122)}$** | -7.7 | 2.9 | -10.6 | 2.27 | 1.7 |

Absolute errors: 0.1-0.2 kcal/mol for $\Delta G$, 0.3-0.5 kcal/mol for $\Delta H$ and $-T\Delta S$, 0.2 for $n$.
Relative error: 20-30% for $K_D$.

**Table S2. Crystallographic data collection and refinement statistics***

| | **C*c*:NPM$_{(9-122)}$ complex** |
|---|---|
| **Data collection** | |
| Wavelength (Å) | 0.97946 |
| Space group | $P2_1$ |
| Unit cell *a, b, c* (Å) | 58.8, 179.72, 104.3 |
| Unit cell β (º) | 94.32 |
| T (K) | 100 |
| X-ray source | Synchrotron |
| Resolution range (Å) | 45.53-(2.55-2.61) |
| Unique reflections | 69833 |
| Completeness (%) | 99.6 (99.8) |
| Redundancy | 6.3 (6.4) |
| $R_{merge}$ | 0.12 (1.22) |
| $R_{pim}$ | 0.08 (0.74) |
| Average $I/\sigma(I)$ | 9.6 (2.1) |
| **Refinement** | |
| Resolution range (Å) | 45.52-2.55 |
| $R_{work}/R_{free}$ | 0.16/0.23 |
| No. Atoms | |
| Protein | 16213 |
| Water | 235 |
| $Cl^-$ | 4 |
| B-factor (Å$^2$) | |
| Protein | 56.93 |
| Water | 47.75 |
| R.m.s. deviations | |
| Bond length (Å) | 0.009 |
| Bond angles (°) | 1.27 |
| Ramachandran | |
| Favored/outliers (%) | 93.00/1.9 |
| Monomers per AU | 20 |
| PDB code | 5EHD |

*Values between parentheses correspond to the highest resolution shells
